# Cdk5 and GSK3β inhibit Fast Endophilin-Mediated Endocytosis

**DOI:** 10.1101/2020.04.11.036863

**Authors:** Antonio P. A. Ferreira, Alessandra Casamento, Sara Carrillo Roas, James Panambalana, Shaan Subramaniam, Kira Schützenhofer, Els F. Halff, Laura Chan Wah Hak, Kieran McGourty, Josef T. Kittler, Konstantinos Thalassinos, Denis Martinvalet, Emmanuel Boucrot

## Abstract

Endocytosis mediates the cellular uptake of micronutrients and cell surface proteins. Parallel to Clathrin-mediated endocytosis, additional Clathrin-independent endocytic routes exist, including fast Endophilin-mediated endocytosis (FEME). The latter is not constitutively active but requires the activation of selected receptors. In cell culture, however, the high levels of growth factors in the regular culture media induce spontaneous FEME, which can be suppressed upon serum starvation. Thus, we predicted a role for protein kinases in this growth factor receptor-mediated regulation of the pathway. Using chemical and genetic inhibition, we found that Cdk5 and GSK3β are negative regulators of FEME. Their inhibition was sufficient to activate FEME promptly in resting cells and boosted the production of endocytic carriers containing β 1-adrenergic receptor, following dobutamine addition. We established that the kinases suppress FEME at several levels. They control Dynamin-1 and Dynein recruitment and sorting of cargo receptors such as Plexin A1 and ROBO1 into FEME carriers. They do so by antagonizing the binding of Endophilin to Dynamin-1 as well as to Collapsin response mediator protein 4 (CRMP4), a Plexin A1 adaptor. Cdk5 and GSK3β also hamper the binding and recruitment of Dynein onto FEME carriers by Bin1. Interestingly, we found that GSK3β binds to Endophilin, thus imposing a local regulation of FEME. Collectively, these findings place the two kinases as key regulators of FEME, licensing cells for rapid uptake by the pathway only upon when Cdk5 and GSK3β activity is low.

---

Clathrin-mediated endocytosis (CME) is the major uptake pathway in resting cells^1,2^ but additional Clathrin-independent endocytic (CIE) routes, including fast Endophilin-mediated endocytosis (FEME), perform specific functions or internalize various cargoes^3,4^. FEME is not constitutively active but is triggered upon the stimulation of selected cell surface receptors by their ligands^**5**^. These include G-protein coupled receptors (*e.g*. β1-adrenergic receptor, hereafter β1AR), receptor tyrosine kinases (*e.g*. epidermal growth factor receptor, EGFR) or cytokine receptors (*e.g*. Interleukin-2 receptor)^5^. In resting cells, FEME is primed by a cascade of molecular events starting with active, GTP-loaded, Cdc42 recruiting CIP4/FBP17 that engage the 5’-phosphatase SHIP2 and Lamellipodin (Lpd). The latter then concentrates Endophilin into clusters on discrete locations of the plasma membrane^6^. In absence of receptor activation, the clusters dissemble quickly (after 5 to 15 sec) upon local recruitment of the Cdc42 GTPase-activating proteins RICH1, SH3BP1 or Oligophrenin^6^ New priming cycles start nearby, constantly priming the plasma membrane for FEME. Upon activation, receptors are quickly sorted into pre-existing Endophilin clusters that then bud to form FEME carriers, which are Clathrin-negative, Endophilin-positive assemblies (EPAs) found in the cytosol. The entire process takes 4 to 10sec ^5^. These FEME carriers travel rapidly to fuse with early endosomes and deliver their cargoes^5^.

Some cell types display spontaneous FEME when grown in their regular culture medium, while others do not. Normal RPE1 cells, primary human dermal fibroblasts (hDFA) and human umbilical vein endothelial cells (HUVEC) exhibited robust FEME in resting cultures grown in regular media (∼5 to 15 EPAs per 100 μm^2^, **Figure S1a-c**). In contrast, HeLa, HEK293 and BSC1 cells displayed very little spontaneous FEME. In these cells, not was FEME identified in a minority of cells but, a low number of FEME carriers were detected in those that were active (∼1 to 3 EPAs per 100 μm^2^, near the leading edge, **Figure S1a-c**). This is not to be confused with FEME priming events (Endophilin short-lived clustering without subsequent carrier budding), which is identified by the growing and disappearance of Endophilin spots without any lateral movements (blinking) by live-cell microscopy. For example, BSC1 cells display abundant priming (clustering of Endophilin) but very little spontaneous FEME (fast moving EPAs into the cytosol)^5,6^. However, within a same culture, not all the cells displayed FEME (the maximum was ∼60% of HUVEC cells, **Figure S1a-b**).

In all cell types tested, FEME was triggered by the addition of 10% serum to complete medium, as shown by an increase in the percentage of cells showing FEME activity as well as increase in EPA production (**Figure S1b-c**). Because FEME was inactive in cells starved of serum^5^ (*i.e. without* growth factors), we looked for kinases that may regulate its activity. Other endocytic pathways are regulated both positively and negatively by multiple protein kinases^7-10^. Thus, we investigated whether phosphorylation would trigger or hinder FEME and looked for a mechanism that may control whether a cell is competent to do FEME or not.

## Results

### Inhibition of Cdk5 and GSK3 activates FEME

A screen of kinases known to regulate membrane trafficking and actin cytoskeleton dynamics (which is required for FEME) was performed using small molecule inhibitors. The compounds used were amongst the best-reported inhibitors for each kinase^11,12^. Four concentrations were tested (10nM, 100nM, 1μM and 10μM), with the minimum effective concentrations for each compound selected for further measurements. Small compounds were chosen because they can be used for short timeframes (minutes), limiting indirect effects on other kinases and long-term cumulative trafficking defects induced by gene depletion. The cells were treated for 10 min at 37°C with the inhibitors diluted in regular growth medium containing 10% serum, and then fixed with pre-warmed paraformaldehyde solution (to preserve FEME carriers, see **Methods**). RPE1 cells were used because they display robust spontaneous FEME when grown in regular medium (**Figure 1a**, ‘normal’). This allowed identification of kinase inhibitors that either decreased or increased FEME. Normal FEME activity (that is, similar to DMSO-treated cells) was given one mark during scoring (**Figure 1a**). Positive and negative controls (dobutamine and PI3Ki, respectively^5^) benchmarked the scoring for ‘decreased’ (zero mark) and ‘increased’ (two marks) FEME (**Figure 1a**). ‘Decreased’ FEME was assigned for samples with >80% reduction in the number of EPAs, in at least 50% of the cells. ‘Increased’ FEME was attributed to samples with >200% elevation in the number of EPAs, in at least 50% of the cells.

These stringent criteria likely missed mild modulations but revealed robust regulators of FEME. Inhibition of CaMKK1 and 2, SYK, FAK or mTORC1/2 reduced FEME significantly (**Figure 1b**), but this was not investigated further in the present study. Conversely, acute inhibition of Cdk5, GSK3 or p38 increased spontaneous FEME (**Figure 1b**). Even though the three inhibitors for Cdk5 also inhibited other cyclin-dependent kinases, the role of the former was deduced by the absence of FEME activation by compounds blocking Cdk1 and 2 (**Figure 1b**), and by the nuclear functions of Cdk7 and 9 ^13^. The role for p38 in FEME was not investigated further at his stage. Cdk5 inhibitors Dinaciclib, Roscovitine and GSK3 inhibitors BIO and CHIR-99021 were validated to activate FEME in a dose-dependent manner (**Figure 1c**) and confirmed that these kinases inhibit FEME in resting cells. Inhibition of Cdk5 or GSK3 increased productive FEME, as a larger number of endocytic carriers contained β1AR upon its activation by dobutamine (**Figure 1d**).

**Figure 1.**
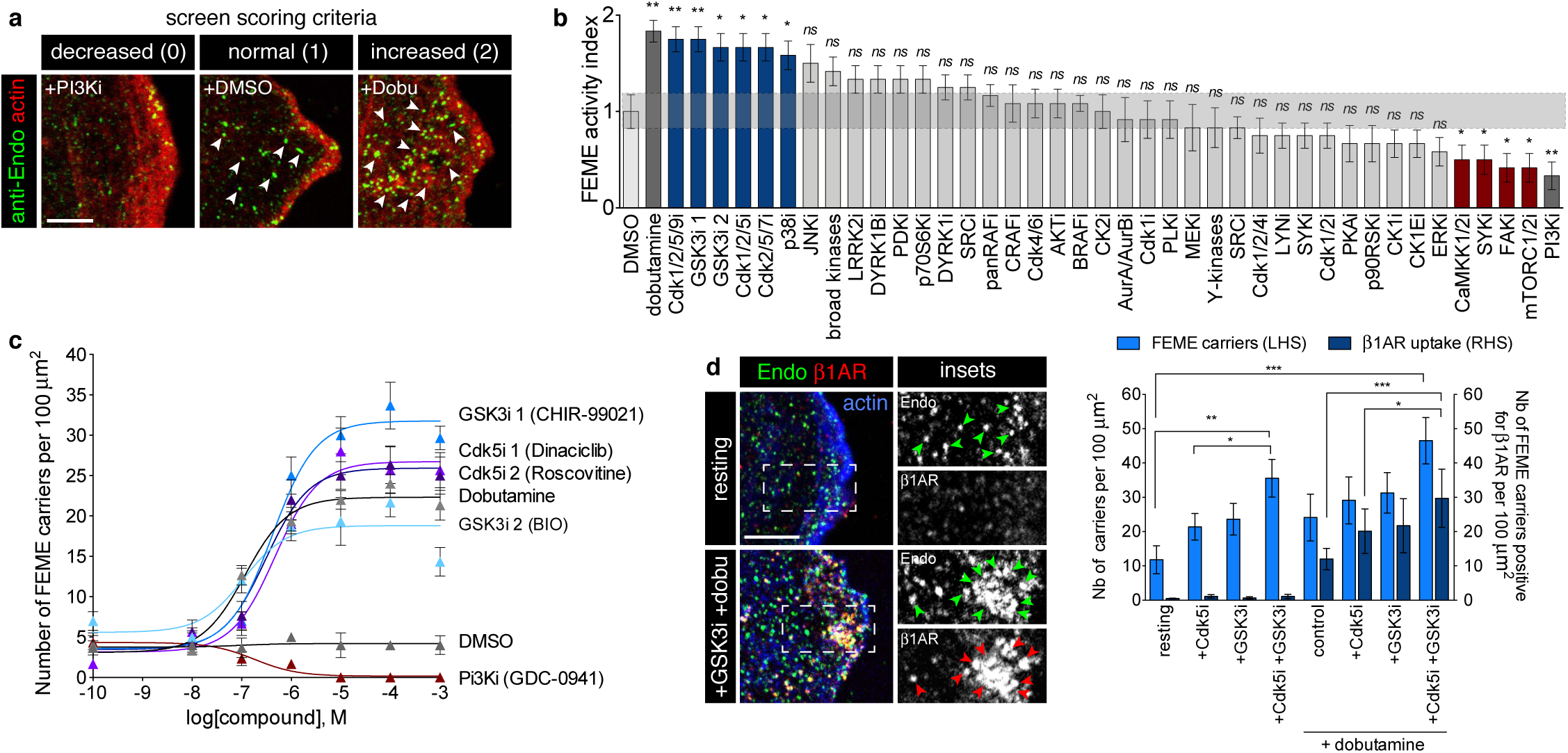
Acute inhibition of Cdk5 and GSK3 activates FEME. **a**, Scoring criteria used in the kinase screen. Representative images of ‘decreased’, ‘normal’ and ‘increased’ FEME in resting human RPE1 cells treated with 10μM dobutamine, 10μM DMSO and 10 nM GDC-0941 (PI3Ki), respectively. Arrowheads point at FEME carriers. ‘Decreased’ FEME was assigned for samples with >80% reduction in the number of EPAs, in at least 50% of the cells. ‘Increased’ FEME was attributed to samples with >200% elevation in the number of EPAs, in at least 50% of the cells. The corresponding scoring marks were 0, 1 and 2, respectively. **b**, Kinase screen using small compound inhibitors. RPE1 cells grown in complete medium were incubated for 10min at 37°C with the following inhibitors: DMSO, (vehicle); dobutamine, 10μM (positive control); Dinaciclib (Cdk1/2/5/9i), 1μM; CHIR-99041 (GSK3i1), 1μM; BIO (GSK3i2), 1μM; Roscovitine (Cdk1/2/5i), 1 m M; PHA-793887 (Cdk2/5/7i), 100nM; VX-745 (p38i), 10μM; JNK-IN-8 (JNKi), 1μM; staurosporine (broad kinases), 1μM; GNE-7915 (LRRK2i), 1μM; GSK2334470 (PDKi), 10μM; PF-4708671 (p70S6Ki), 10μM; AZ191 (DYRKi), 10μM; AZD0530 (SRCi), 1μM; TAK-632 (panRAFi), 10μM; GW 5074 (CRAFi), 1μM; PD0332991 (Cdk4/6i), 1μM; MK2206 (AKTi), 1μM; GDC-0879 (BRAFi), 1μM; CX-4945 (CK2i), 1μM; ZM 447439 (AurA/AurBi), 1μM; RO-3306 (Cdk1i), 100nM; BI 2536 (PLKi), 1μM; PD0325901 (MEKi), 100nM; Genistein (Y-kinases), 1μM; Purvalanol A (Cdk1/2/4i), 100nM; MLR 1023 (LYNi), 1μM; CDK1/2 inhibitor III (Cdk1/2i), 100nM; KT 5720 (PKAi), 100nM; BI-D1870 (p90RSKi), 100nM; PF-4800567 (CK1Ei), 1μM; SCH772984 (ERKi), 100nM; STO609 (CaMKK1/2ii), 100nM; P505-15 (SYKi), 1μM; PND-1186 (FAKi), 100nM; Torin 1 (mTORC1/2i), 10μM and GDC-0941 (PI3Ki), 100nM (negative control). **c**, Number of FEME carriers (cytoplasmic Endophilin-positive assemblies, EPAs) upon titration of CHIR-99021, BIO, Roscovitine and Dinaciclib. Dobutamine and GDC-0941 were used as positive and negative controls, respectively. **d**, β1-adrenergic receptor (β1AR) uptake into FEME carriers in RPE1 cells pre-treated with 5μM CHIR-99021 (GSK3i) for 5 min, followed by 10μM dobutamine for 4 min or not (resting). Histograms show the mean ± SEM of the number of FEME carriers (left axis) and the number of FEME carriers positive for β1AR per 100 μm^2^ (right axis) (*n*=30 cells per condition, from biological triplicates). Arrowheads point at FEME carriers. All experiments were repeated at least three times with similar results. Statistical analysis was performed by one-way ANOVA (b, and c) or two-way ANOVA (d); *NS*, non significant; *, *P*<0.05, **, *P* <0.01, ***, *P* <0.001. Scale bars, 5μm.

### Endophilin recruits GSK3β to regulate FEME locally

GSK3α and β kinases require prior phosphorylation of its substrates by several other kinases (*e.g*. PKA, AMPK, CK1/2, Cdk5). GSK3 docks onto the priming sites and then phosphorylates nearby Serines or Threonines, a few amino acids away^14^. The kinase is auto-inhibited by phosphorylation at its N-terminus (Ser 9 in GSK3β, pS9-GSK3β hereafter), which then occupies its docking site and thus blocks its interaction with substrates. This phosphorylation on GSK3 is mediated by many kinases that are activated by growth factor receptor signaling including AKT and ERK^15,16^. In absence of growth factors (such as upon serum starvation), GSK3β phosphorylation on Ser 9 is reduced, relieving auto-inhibition of the kinase^15^. Consistently, we observed reduced levels of inactive pS9-GSK3β and depression of FEME in cells starved for growth factors (**Figure 2a-b**). In contrast, stimulating cells with an additional 10% serum (20% final) for 10 min inactivated the kinase (as deduced from the high levels of inactive pS9-GSK3β), and activated FEME beyond resting levels (**Figure 2a-b**). There was a linear correlation (r^2^=0.99) between the levels of inactive GSK3β and the numbers of FEME carriers within the same cells (**Figure 2a-b**). However, activation of other receptors (*e.g*. β1AR with dobutamine) activated FEME in resting cells without measurable changes in GSK3β activity (**Figure 2b**, ‘+dobutamine’, middle). There was a maximum level of FEME activity, as addition of dobutamine on cells that were previously activated with 10% extra serum did not increase the number of EPAs further (**Figure 2b**, ‘+dobutamine’, right). Interestingly, when GSK3β activity was high (starved cells), dobutamine could not activate FEME (**Figure 2b**, ‘+dobutamine’, left). There was a poor correlation (r^2^=0.67) between the kinase activity and FEME stimulation by dobutamine, suggesting that GSK3β acts upstream, imposing a cap on FEME that can be quickly lifted upon receptor activation.

**Figure 2.**
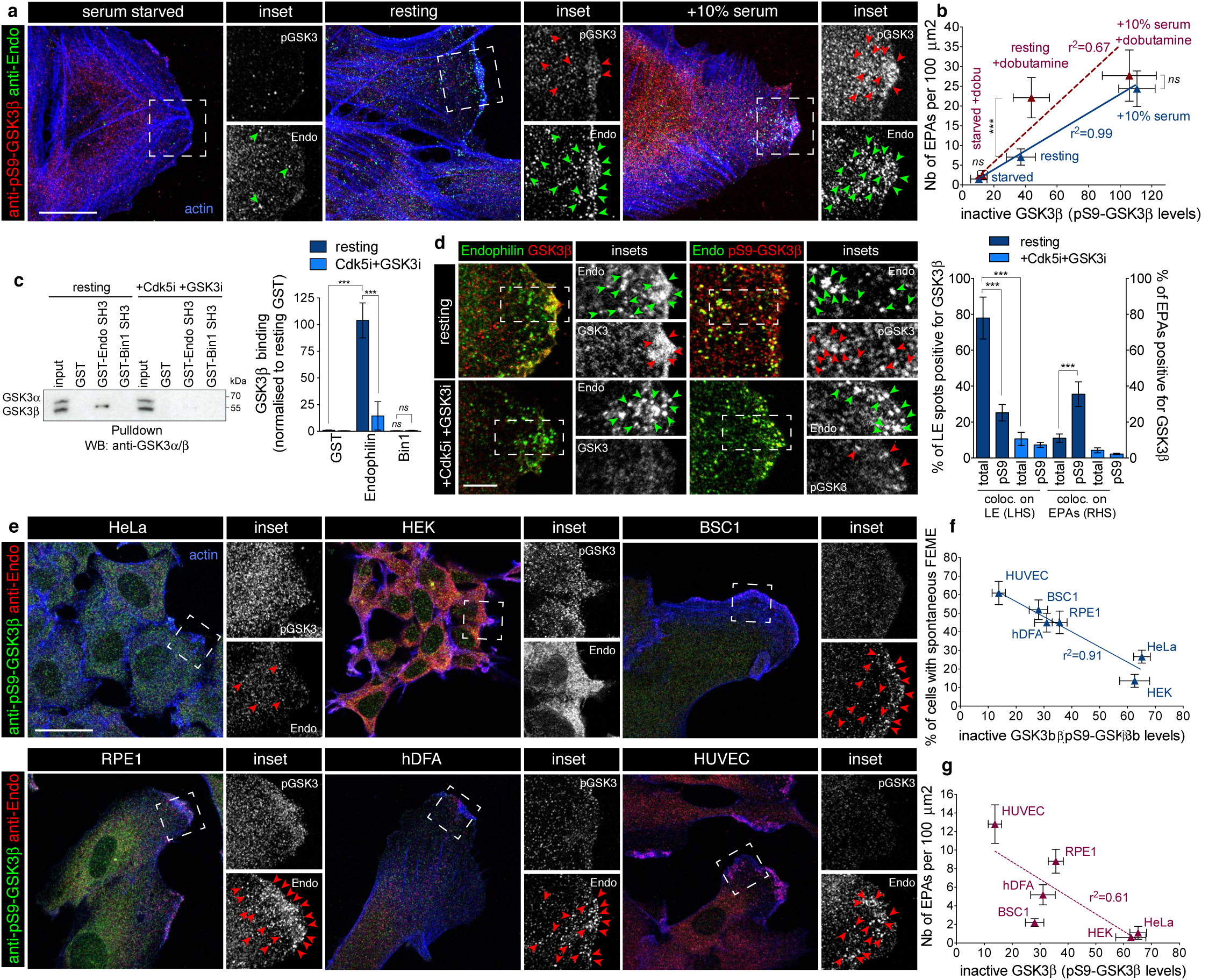
Endophilin recruits GSK3 b for local regulation of FEME. **a**, Confocal images showing levels of phosphorylated Ser9 GSK3β (inactive kinase) and colocalization with Endophilin in cells starved of serum for 1h (‘serum starved’), grown in 10% serum medium (‘resting’) or stimulated with additional serum for 10 min (‘+10% serum’)). Arrowheads point at FEME carriers. **b**, Correlation between the number of EPAs and pS9-GSK3β levels (single cell measurements) in cells that were starved of serum for 1h (‘starved’), grown in 10% serum medium (‘resting’) or stimulated with additional serum for 10 min (‘+10% serum’), followed by the addition of 10 m M dobutamine for 4 min (red data points) or not (blue data points). **c**, Left, pull-down experiments using beads with GST-SH3 domains of Endophilin A2 or Bin1, in resting cells or cells treated with 5μM Dinaciclib (Cdk5i) and CHIR-99021 (GSK3i) for 10 min. GST beads were used as negative control. Bound GSK3β was detected using an antibody that detects both GSK-3*α* and GSK-3β. Right, histograms show the mean ± SEM of GSK3β binding, normalized to resting GST levels. **d**, Colocalization of total and phosphorylated Ser9 (inactive) GSK3β and Endophilin in cells treated with 5μM Cdk5 and GSK3 inhibitors for 10 min, or not (resting). Histograms show the mean ± SEM of Endophilin spots at the leading edge of cells (spots within 1 m m of cell edges) and on EPAs positive for total or pS9-GSK3β. (*n*=50 spots or EPAs per condition, from biological triplicates). Arrowheads point at Endophilin spots and FEME carriers. **e**, Confocal images showing levels of phosphorylated Ser9 GSK3β (inactive kinase) and colocalization with Endophilin in resting HeLa, HEK, BSC1, RPE1, hDFA or HUVEC grown in their respective full serum media. Arrowheads point at FEME carriers. **f**, Correlation between the percentage of cells displaying active FEME and their pS9-GSK3β levels (single cell measurements) in the indicated resting cell types. **g**, Correlation between the number of EPAs and pS9-GSK3β levels (single cell measurements) in the indicated resting cell types. All experiments were repeated at least three times with similar results. Linear regression fit (or absence thereof) is indicated as r^2^ values. Statistical analysis was performed by one-way ANOVA (b, c, f and g); *NS*, non significant, *, *P*<0.05, **, *P* <0.01, ***, *P* <0.001. Scale bars, 20 (a and e) and 5μm (d).

As mass spectrometry detected GSK3β amongst the proteins immunopurified by anti-Endophilin antibodies (**Supplementary Figure 2a and Table 1**), we further characterized its binding to Endophilin. We found that GSK3β, but not α, bound to the SH3 domain of Endophilin but not that of the closely related N-BAR domain protein Bin1 (**Figure 2c**). Consistently with the detection of GSK3β from resting but not FBS-stimulated extracts, (**Supplementary Figure 2a**), the binding was not detected in extracts of cells that were inhibited for Cdk5 and GSK3 prior to lysis (**Figure 2c**). Furthermore, GSK3β was detected on priming Endophilin spots at the leading edge but little on FEME carriers following inhibition (**Figure 2d**). In contrast, inactive GSK3β was more detected on EPAs than on priming spots at the leading edge (**Figure 2d**). Consistent with binding data, Cdk5 and GSK3 inhibitors blocked the recruitment of both total and inactive GSK3β (**Figure 2d**). Thus, we concluded that Endophilin recruits GSK3β to inhibit FEME locally. However, we found that levels of the inactive pS9-GSK3β were inversely correlated with the proportion of resting cells having spontaneous FEME (r^2^=0.91, **Figure 2e-f**). But there was no correlation between the kinase activity and the amounts of EPAs produced by spontaneous FEME in these resting cells (r^2^=0.61, **Figure 2g**). Because dobutamine did activate FEME robustly, regardless of basal GSK3β activity, we hypothesized that there must be another layer of regulation upstream or parallel to the kinase. Given the requirement of a priming kinase for GSK3 action^17,14^, we tested a potential synergy with Cdk5.

### Cdk5 and GSK3β work in synergy to inhibit FEME

Genetic inhibition of either Cdk5 or GSK3α/β increased spontaneous FEME in resting cells (**Figure 3a-b** and **Supplementary Figure S2b**), and, conversely, in individual cells, the levels of the kinases correlated negatively with FEME activity (r^2^=0.97, **Figure 3b**). This confirmed the data obtained with the kinase inhibitors, but also suggests that as long as the kinases are inactive, FEME is elevated, as gene depletion by RNAi lasts several days. RNAi rescue with high levels of either wild type (WT) or constitutively active (CA) forms of the kinases were sufficient to suppress FEME (**Figure 3a, c**). However, dominant-negative forms of either Cdk5 or GSK3β did not rescue the effect of the depletion of the endogenous kinases. The dual inhibition of Cdk5 and GSK3β occasioned a synergistic activation of FEME (**Figure 3d-e and 1d**), driving not only cargo loading but also FEME carrier budding and lateral movement within the cytoplasm.

Dynamin mediates the budding of FEME carriers^5^ and is known to be phosphorylated on Ser778 by Cdk5 ^18^, and subsequently on Ser774 by GSK3β ^9^ (**Figure 4a)**. This blocks the function of Dynamin in CME but is required for activity-dependent bulk endocytosis in synapses^8,9^. Consistent with proximity of the phosphorylated residues to the binding site of Endophilin on Dynamin^19,20^, the inhibition of both Cdk5 and GSK3 increased Dynamin recruitment onto budding FEME carriers (**Figure 3e and 4a-d**). As the single inhibition of either Cdk5 or GSK3β was sufficient to relieve FEME from their control (even though the other kinase should not be affected), we concluded that Cdk5 acts upstream of GSK3β, and that other kinases may prime GSK3β in absence of Cdk5. The synergy of their inhibition suggested that they hamper FEME at several steps, beyond Dynamin recruitment.

**Figure 3.**
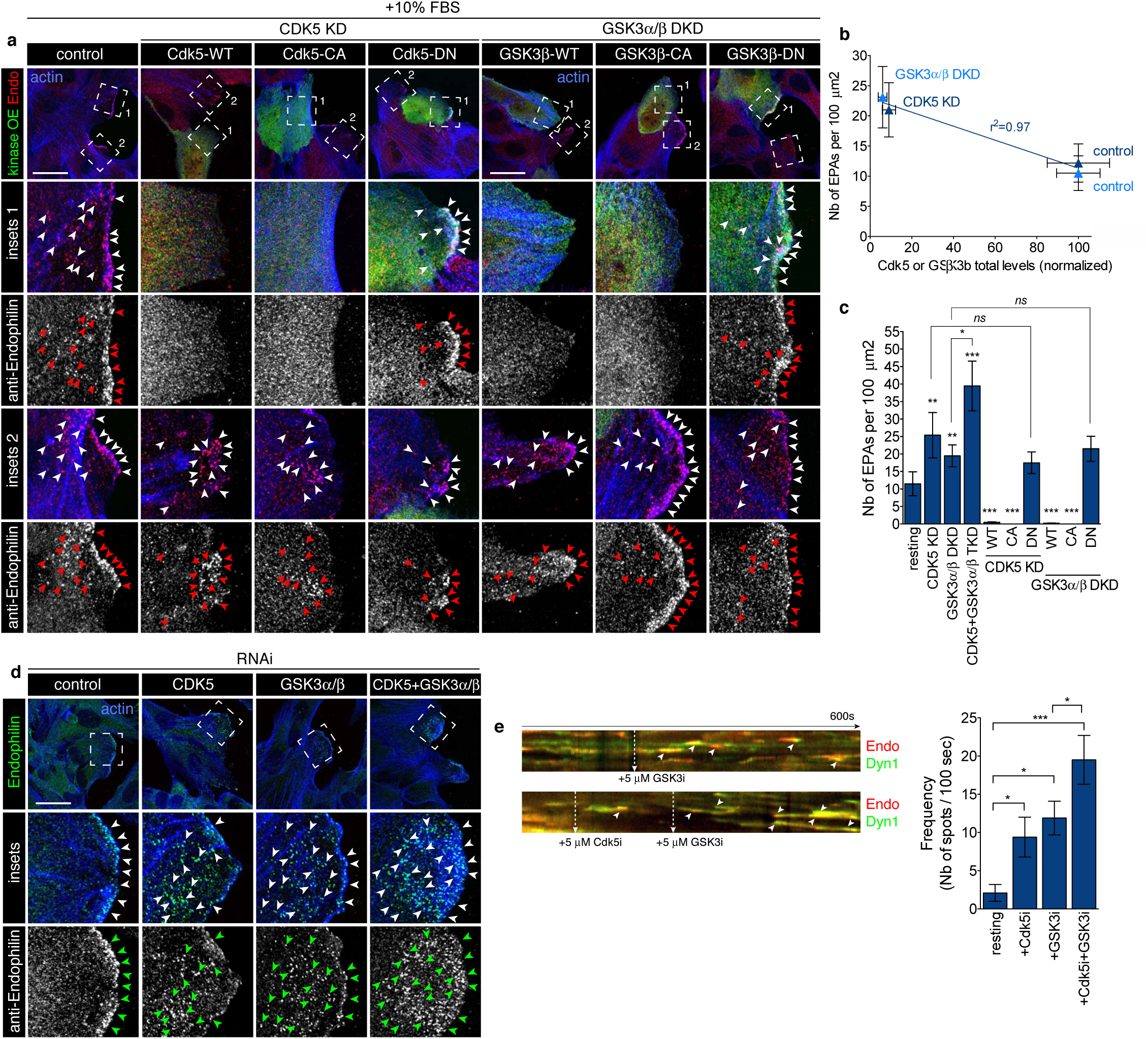
Cdk5 and GSK3β act in synergy to control FEME. **a**, Confocal microscopy images of control RPE1 cells (‘control), or cells in which Cdk5 or GSK3α and β had been knocked-down using RNAi (‘CDK5 KD’ or ‘GSK3α*/*β DKD’, respectively) cells. Wild-type (WT), constitutively active (CA) or dominant negative (DN) forms of Cdk5 or GSK3β (green) were overexpressed in a knock-down background and endogenous Endophilin (red) was immunostained, as indicated. All cells were stimulated with +10% FBS (20% final) for 10min prior to fixation. Arrowheads point at FEME carriers. **b**, Correlation between the number of EPAs and their Cdk5 or GSK3β levels (single cell measurements) in control of Cdk5 or GSK3α/β depleted cells, as indicated. **c**, Number of EPAs in RPE1 cells treated as indicated in a and d. **d**, Confocal microscopy images of resting control RPE1 cells (‘control), Cdk5 (‘CDK5 KD), GSK3α and β (‘GSK3α/β DKD’) or Cdk5 and GSK3α/β (‘Cdk5+GSK3α/β TKD’) knocked-down cells. Arrowheads point at FEME carriers. **e**, Kymographs from cells expressing low levels of Dynamin2-EGFP and EndophilinA2-RFP, treated with CHIR-99021 (GSK3i) or Dinaciclib (Cdk5i) as indicated and imaged at 2Hz. Arrowheads point at FEME carriers. Kymographs are representative of at least 3 captures from biological triplicates. Histograms show the mean ± SEM from biological triplicates (*n*=3 cells per condition). All experiments were repeated at least three times with similar results. Linear regression fit is indicated as r^2^ value. Statistical analysis was performed by one-way ANOVA (b, c and e); *NS*, non significant; *, *P*<0.05, **, *P* <0.01, ***, *P* <0.001. Scale bars, 20 μm.

**Figure 4.**
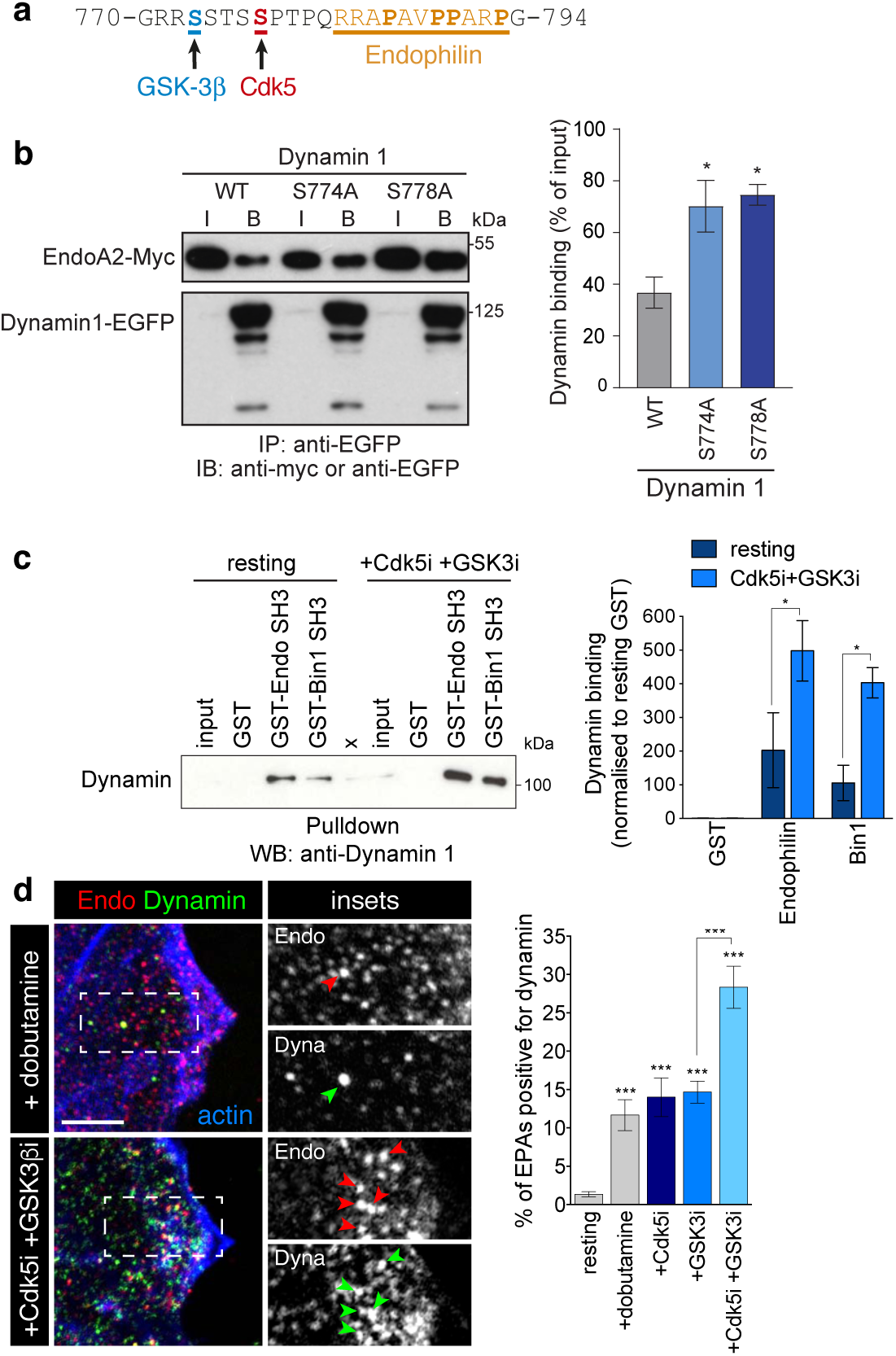
Cdk5 and GSK3β regulate Dynamin recruitment onto FEME carriers. **a**, Human Dynamin-1 sequence (aa 770-794). Amino acids phosphorylated by GSK3β (S774) and by Cdk5 (S778) are shown in blue and red, respectively. The proline-rich motif to which Endophilin is known to bind to is shown in orange (underlined). **b**, Co-immunoprecipitation of Endophilin A2-Myc and Dynamin1-EGFP wild-type (WT), or non-phosphorylatable mutants S774A or S778A. Inputs (I) correspond to 0.5% of the cell extracts), and bound fractions (B) to 90% of material immunoprecipitated. Righ, Histograms show the mean ± SEM from three independent biological experiments. **c**, Pull-down experiments using beads with GST-SH3 domains of Endophilin A2 or Bin1, in resting cells or cells treated with 5μM Dinaciclib (Cdk5i) and CHIR-99021 (GSK3i) for 10 min. GST beads were used as negative control. ‘X’ labels a lane that was not used in this study. Inputs correspond to 4% of cell extracts. Right, histograms show the mean ± SEM of Dynamin-1 binding, normalized to resting GST levels. **d**, Recruitment of endogenous Dynamin onto FEME carriers in RPE1 cells treated for 10 min with 5μM Dinaciclib (Cdk5i) and/or CHIR-99021 (GSK3i) or not, followed by 10μM dobutamine for 4 min. Arrowheads point at FEME carriers. Histograms show the mean ± SEM from biological triplicates (*n*=50 cells per condition). All experiments were repeated at least three times with similar results. Statistical analysis was performed by one-way ANOVA (b and d) or two-way ANOVA (c); *NS*, non significant; *, *P*<0.05, **, *P* <0.01, ***, *P* <0.001. Scale bar, 5μm.

### Cdk5 blocks the binding of Endophilin to CRMP4 and the sorting of PlexinA1 into FEME carriers

Amongst proteins known to be phosphorylated by Cdk5 and GSK3β is collapsin response mediator protein 4 (CRMP4)^21^. Interestingly, we recently identified CRMP1, 3 and 4 in pulldown experiments using Endophilin SH3 domains^6^. CRMP1 to 5 form homo and hetero tetramers acting as adaptors during cell guidance mediated by Plexin A1 (**Figure 5a)**, and mediate cytoskeletal remodeling upon Semaphorin 3A or 6D sensing^22-24^. Cdk5 phosphorylates CRMP4 at Ser 522, which primes GSK3β-mediated phosphorylation at the positions Thr 509, Thr 514 and Ser 518 ^21^ (**Figure 5a**). The phosphorylation of CRMP4 perturbs its binding to microtubules and actin and it is critical for proper neuronal development in both zebrafish and mice^25,26^. Inhibiting Cdk5, or overexpressing the non-phosphorylatable mutant CRMP4-S522A, increased CRMP4 binding to Endophilin, whereas a phospho-mimetic mutation S522D had the opposite effect (**Figure 5b-c** and **Supplementary S3a-b**). Point mutations P526A or R525E, but not P502A, abolished the interaction (**Figure 5d** and **Supplementary S3c**), establishing that the binding motif for Endophilin on CRMP4 is the proline-rich motif (aa 523-529) proximal to Ser 522 (**Figure 5a**). The sites phosphorylated by GSK3 are several amino acids away from that motif, explaining why GSK3 inhibition did not affect the interaction of Endophilin with CRMP4 (**Figure 5b** and **Supplementary S3a**). Consistent with the biochemical data, mutations in CRMP4 inhibiting its binding to Endophilin abrogated localization of CRMP4 on FEME carriers, even upon co-overexpression (**Figure 5e** and **Supplementary S3d**). In HUVEC cells, which express CRMP4 and Plexin A1 endogenously^27^,Cdk5 inhibition enhanced the recruitment of endogenous CRMP4 onto FEME carriers and the uptake of Plexin A1 upon Semaphorin 3A stimulation (**Figure 5f-g** and **Supplementary S3e-f**). Similarly to other receptors (*e.g*. EGFR), PlexinA1 used both FEME and CME to enter cells **(Supplementary S3e**), perhaps from different cellular location (at the leading edge of cells, most PlexinA1 internalized into FEME carriers (**Figure 5g** and **Supplementary S3f**). Altogether, this established that Cdk5 blocks the binding of Endophilin to CRMP4 and uptake of Plexin A1 in FEME carriers.

**Figure 5.**
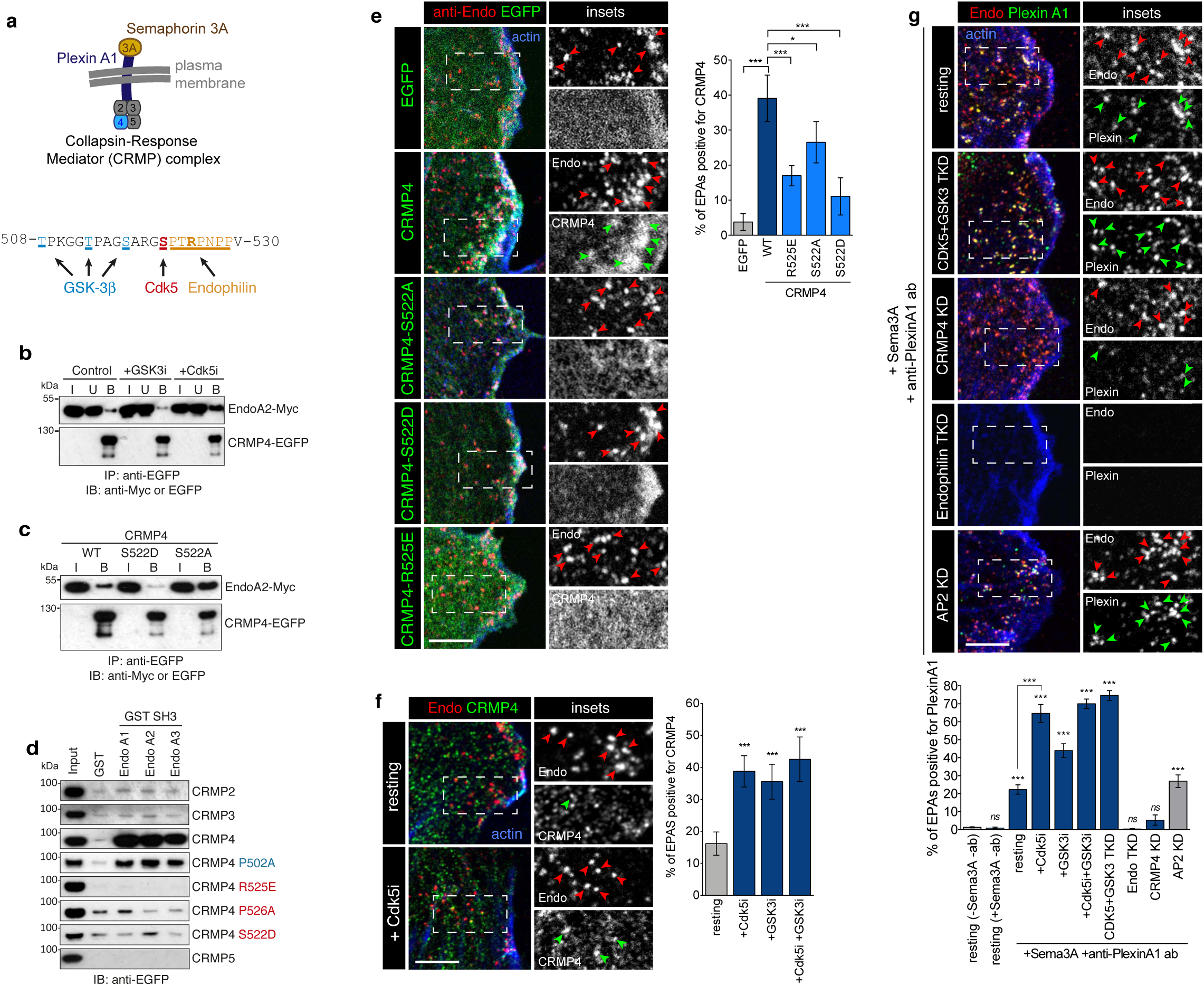
Cdk5-mediated phosphorylation of CRMP4 inhibits the binding of Endophilin to CRMP4 and the sorting of Plexin A1 into FEME carriers. **a**, Top, diagram showing the recruitment of CRMP2-5 adaptor complex (CRMP4 highlighted in blue) to Plexin A1 upon stimulation with Semaphorin 3A. Bottom, human CRMP4 protein sequence (aa 508-530). Amino acids phosphorylated by GSK3β (T509, T514 and S518) and by Cdk5 (S522) are shown in blue and red, respectively. The Endophilin binding motif established in this study is shown in orange (underlined). **b**, co-immunoprecipitation of CRMP4-EGFP and Endophilin A2-Myc from cells treated with 5μM CHIR-99021 (GSK3i) or Dinaciclib (Cdk5i) as indicated. I, input (10% of the cell extracts), U, unbound (10% of total) and B, bound fractions (80% of total), respectively. **c**, co-immunoprecipitation of CRMP4-EGFP wild-type (WT), S22D or S522A and Endophilin A2-Myc. I, input (10% of the cell extracts), and B, bound fractions (90% of total), respectively. **d**, Pull-down using GST-SH3 domains of Endophilin A1, A2 or A3 and cell extracts expressing the indicated EGFP-tagged CRMP proteins. GST was used as negative control. Binding proteins were detected by immunoblotting with an anti-EGFP antibody. ‘input’ lanes correspond to 5% of the cell extracts. **e**, Recruitment of EGFP-tagged CRMP4 WT, S522D, S522A or R525E onto FEME carriers (cytoplasmic Endophilin-positive assemblies, EPAs) in HUVEC cells. Arrowheads point at FEME carriers. Right, histograms show the mean ± SEM from biological triplicates (*n*=30 cells per condition). **f**, Recruitment of endogenous CRMP4 onto FEME carriers in HUVEC cells treated for 10min with 5μM Dinaciclib (Cdk5i) and/or CHIR-99021 (GSK3i), or not (resting). Arrowheads point at FEME carriers. Histograms show the mean ± SEM from biological triplicates (*n*=45 cells per condition). **g**, Endogenous Plexin A1 uptake into FEME carriers in HUVEC cells depleted of Endophilin A1, A2 and A3 (‘Endophilin TKD), Cdk5 and GSK3α and b (‘CDK5+GSK3α/β TKD’), CRMP4 (‘CRMP4 KD’) or AP2 (‘AP2 KD’) or pre-treated with Cdk5i and/or GSK3i for 5min. Cells were stimulated by 20nM Semaphorin 3A (Sema3A) for 5min in presence of 10 μg/mL anti-PlexinA1 antibodies (recognizing the ectodomain of PlexinA1) or not (resting). Arrowheads point at FEME carriers. Histograms show the mean ± SEM from biological triplicates (*n*=30 cells per condition). All experiments were repeated at least three times with similar results. Statistical analysis was performed by one-way ANOVA; *NS*, non significant; *, *P* <0.05, ***, *P* <0.001. Scale bars, 5μm.

Endophilin recruits cargoes into FEME carriers through the binding of its SH3 domain to proline rich motifs present in cargo adaptors or cytoplasmic tails of receptors^5^. We tested interaction with other cell guidance receptors containing putative proline-rich motifs in their cytoplasmic tails, and found that Endophilin bound to Semaphorin 6A and 6D, and to ROBO1 (**Supplementary S4a**). Roundabout (ROBO) receptors bind to Slit ligands to mediate cell guidance, including axon repulsion^28^. Recently, Endophilin was found to mediate the uptake of ROBO1 and VEGFR2 via a Clathrin-independent pathway reminiscent to FEME^29^. We confirmed that Slit1 enter cells into FEME carriers (**Supplementary S4b**) and its cellular uptake was strongly reduced in FEME-but not in CME-deficient cells (Endophilin triple knock-down ‘TKD’ and AP2 knock-down, respectively, **Supplementary S4b**). Upon binding to Slit, ROBO1 activates AKT, which in turn phosphorylates GSK3β on Ser9, thereby de-activating the kinase in axons^30,31^. Consistently, acute inhibition of GSK3β increased the uptake of Slit1 into FEME carriers two-fold (**Supplementary Figure S5b**). Thus, Cdk5 and GSK3β kinases act at another level of FEME by controlling the sorting of cargoes, such as PlexinA1 and ROBO1, into endocytic carriers.

### Cdk5 and GSK3β controls the recruitment of Dynein by Bin1 onto FEME carriers

As Cdk5 and GSK3 kinases regulate Dynein^32,33^ and because FEME carriers containing Shiga toxin rely on this microtubule motor for scission and retrograde trafficking^34,35^, a role for the kinases in regulating Dynein during FEME was explored. We confirmed that inhibiting Dynein (with either the small inhibitor Ciliobrevin D^36^ or the overexpression of the p50 dynamitin subunit of the dynactin complex^37^) blocked FEME carrier budding and β1AR endocytosis (**Supplementary Figure S5a-b**). However, inhibition of Kinesin upon overexpression of the TPR subunit^38^ had no effect (**Supplementary Figure S5a-b**). Most FEME carriers were detected in the vicinity of microtubules, and their mild depolymerization using low doses (100nM) of nocodazole stalled FEME carriers at the plasma membrane (**Figure 6a**). FEME carriers, produced in resting cells upon acute Cdk5 and GSK3β inhibition, recruited Dynein and travelled along microtubules (**Figure 6a-b**). Dynein was immunopurified together with Endophilin from cell extracts in which Cdk5 was inhibited (**Figure 6c**). To confirm that Dynein was recruited onto FEME carriers produced upon Cdk5 and GSK3β inhibition, immunoprecipitation was performed on membrane fractions enriched in Endophilin but poorer in other endocytic markers (Fractions 7 of sucrose gradients, **Figure 6d**). The material that was immuno-isolated from such fractions likely contained FEME carriers, as they were rich in Endophilin and lipids but devoid of Clathrin. Importantly, Dynein was indeed immunopurified together with Endophilin from such fractions (**Figure 6d-e**).

**Figure 6.**
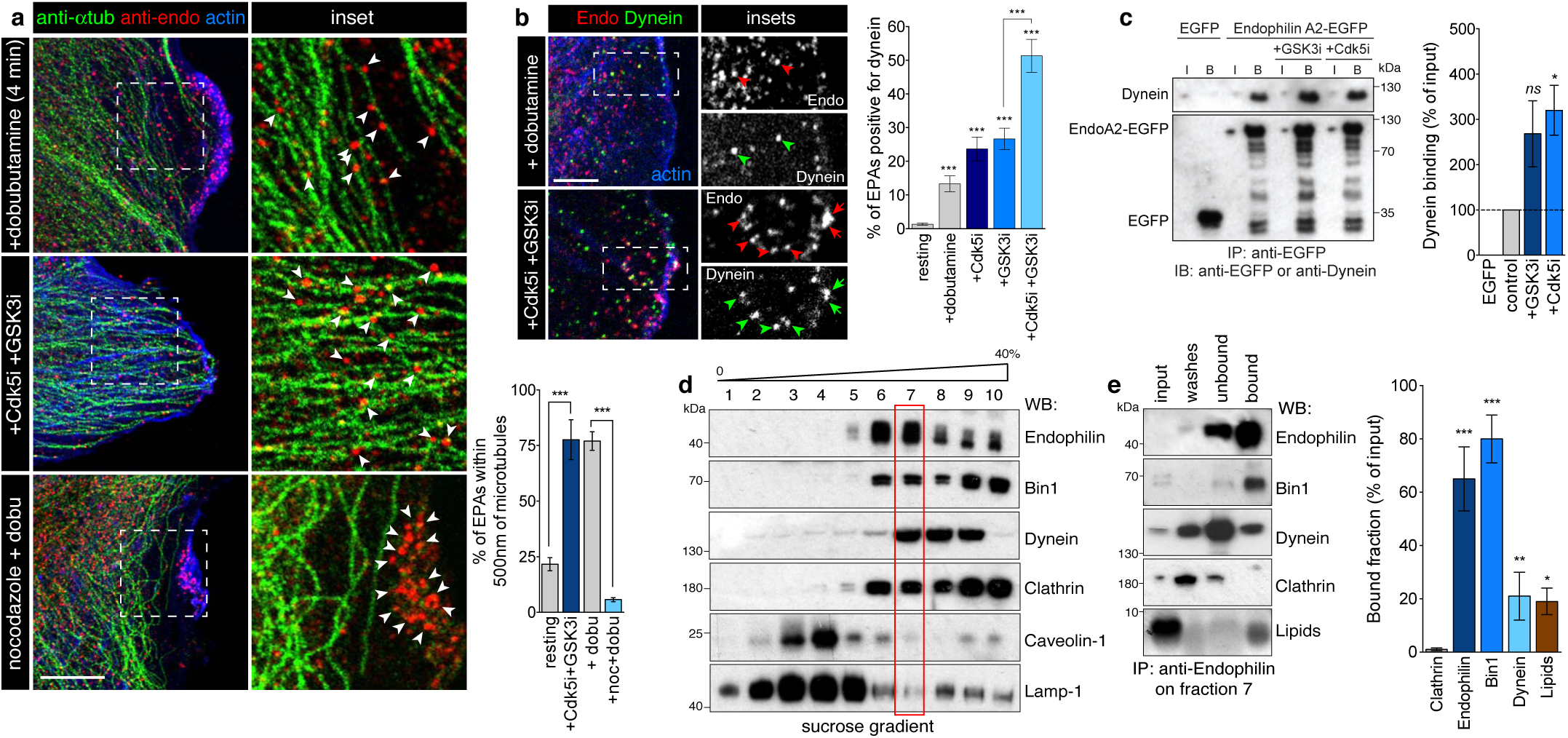
Cdk5 and GSK3β inhibit Dynein recruitment onto FEME carriers. **a**, Juxtaposition of FEME carriers and microtubules in HUVEC cells treated with 10μM dobutamine for 4min, 5μM Dinaciclib (Cdk5i) and CHIR-99021 (GSK3i) for 10min, but not upon mild depolymerization (using 100nM nocodazole for 10min prior to dobutamine stimulation). Arrowheads point at FEME carriers. Right, histograms show the mean ± SEM from biological triplicates (*n*=100 puncta per condition). **b**, Recruitment of endogenous Dynein onto FEME carriers in HUVEC cells treated for 10min with 5μM Dinaciclib (Cdk5i) and CHIR-99021 (GSK3i) or not, followed by 10μM dobutamine for 4min. Arrowheads point at FEME carriers. Histograms show the mean ± SEM from biological triplicates (*n*= 50 cells per condition). **c**, Co-immunoprecipitation experiments with EGFP or Endophilin A2-EGFP, in resting cells or cells treated for 30min with 5μM Dinaciclib (Cdk5i) and CHIR-99021 (GSK3i). I, input (10% of the cell extracts), and B, bound fractions (90% of total), respectively. Righ, Histograms show the mean ± SEM from three independent biological experiments **d**, Sucrose gradient (0 to 40%) membrane isolation form RPE1 cells stimulated with 10%FBS (20% final) for 10min. Fractions were immunoblotted for Endophilin, Bin1, Dynein, Clathrin, Laveolin-1 and Lamp-1. Fraction 7, containing high levels of Endophilin but low levels of Clathrin, Caveolin-1 and Lamp-1 was selected for subsequent immuno-precipitation. **e**, Left, anti-Endophilin immuno-precipitation from fraction 7 samples. Immunoblots measured the levels of Endophilin, Bin1, Dynein, Clathrin and lipids (see Methods) in input (5% of cell extracts), washes (10% of total), unbound (10% of total) and bound (50% of total) samples. Right: Histograms show the mean ± SEM from biological triplicates. .All experiments were repeated at least three times with similar results. Statistical analysis was performed by one-way ANOVA; *NS*, non significant; *, *P* <0.05, **, *P* <0.01, ***, *P* <0.001. Scale bars, 20 (a) and 5 m m (b).

We were intrigued by the presence of Bin1 in the immunoprecipitated fractions, as we initially included it as a control (Bin1 is a N-BAR and SH3 domain-containing protein related to Endophilin). To test for a potential role for Bin1 in FEME, we screened our library containing 72 full-length human BAR proteins tagged with EGFP^6^. But instead of looking for BAR proteins colocalizing onto the transient Endophilin clusters at the leading edge^6^, we focused on those that localized onto FEME carriers produced upon FBS addition (**Figure 7a**). While 10 BAR domain proteins (FAM92B, SH3BP1, ASAP1, SNX9, SNX33, CIP4, Pacsin2, PSTPIP1 and Nostrin) were significantly detected onto a subset (∼15 to 45%) of EPAs, only Amphiphysin, Bin1 and Bin2 located to the majority (>50%) of FEME carriers (**Figure 7a-b**). This partial localization of the 10 aforementioned BAR proteins could be the result of them marking discreet steps and/or sub-population of FEME carriers, or could be simply caused by the ectopic expression of the constructs. These were not studied further at this point. Amongst the three best hits, we focused on Bin1 because it is ubiquitously expressed, unlike Amphiphysin that is brain-enriched^39,40^. Bin2 is a known binding partner of Endophilin that is mainly expressed in leukocytes and that heterodimerizes with Bin1 but not Amphiphysin ^41,42^. Bin1 has several splice variants, including brain-specific long isoforms 1 to 7 (also known as Amphiphysin II) that contain AP2- and Clathrin-binding motifs and function in CME^43^ The two ubiquitously-expressed, short isoforms 9 and 10, however, resemble Endophilin in that they have a N-BAR domain, a short linker and SH3 domain and colocalize poorly with either AP2 or Clathrin (immunostaining, **Supplementary Figure S6a**). Endogenous Bin1 localized onto the majority of FEME carriers produced upon either β1AR activation or Cdk5 and GSK3β inhibition (**Figure 7c**). Like Endophilin^6^, Bin1 binds to Lpd (**Supplementary Figure S6d**) and relies on both CIP4 and Lpd for it recruitment into the transient clusters priming the leading edges of resting cells (**Supplementary Figure S6b-c**). In cells depleted for Bin1 (Amphiphysin and Bin1 double knock-down, ‘Amph+Bin1 DKD’ was performed to avoid potential compensation), CIP4, Lpd and Endophilin were recruited as in control cells (**Supplementary Figure S6b-c**) and β1AR uptake was not affected^6^. This suggested that Bin1 could be mediating the uptake of different cargoes other than β1AR and/or that it could have a later role, post budding. In absence of data supporting or rebutting the first hypothesis, we focused on the potential link with Dynein that we reported above.

**Figure 7.**
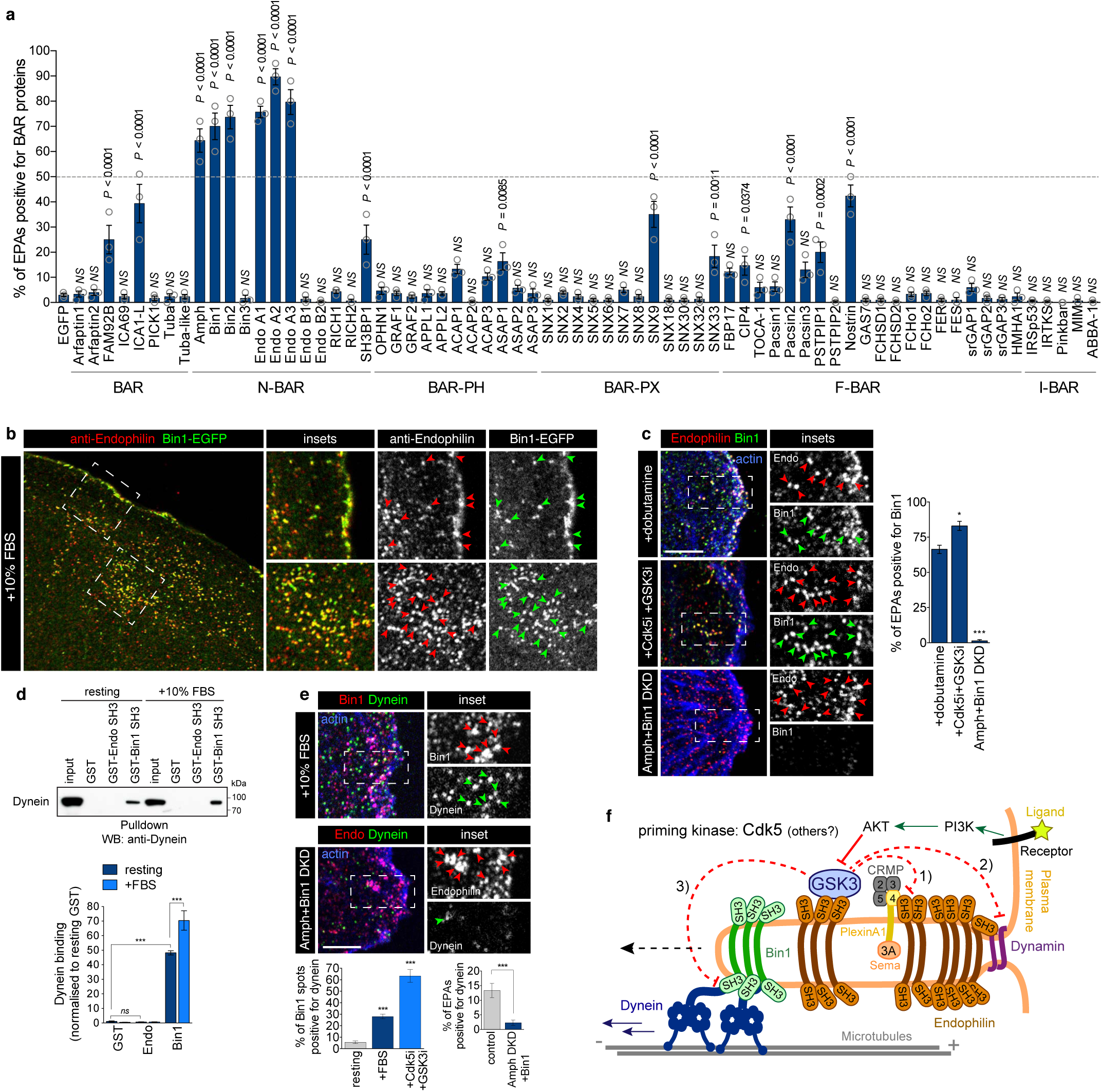
Bin1 recruits Dynein onto FEME carriers. **a**, Colocalization of named EGFP-tagged BAR proteins on FEME carriers marked by endogenous Endophilin in BSC1 cells stimulated with additional 10% serum for 10 min prior to fixation. Histograms show the mean ± SEM from three independent biological experiments (*n*>100 puncta per condition). **b**, Colocalization of Bin1-EGFP on FEME carriers marked by endogenous Endophilin in BSC1 cells stimulated with additional 10% serum for 10 min prior to fixation. Arrowheads point at FEME carriers. **c**, Colocalization of endogenous Bin1 and Endophilin upon stimulation with dobutamine, after treatment with 5μM Dinaciclib (Cdk5i) and CHIR-99021 (GSK3i) for 10min, or in cells depleted of Bin1 Amphiphysin (‘Amph+Bin1 DKD’). Arrowheads point at FEME carriers. Right, histograms show the mean ± SEM from three independent biological experiments (*n*>150 puncta per condition). **d**, Pull-down experiments using beads with GST-SH3 domains of Endophilin A2 or Bin1, in resting cells or cells treated with extra 10% FBS for 10 min. GST beads were used as negative control. Inputs correspond to 4% of cell extracts. Bottom, histograms show the mean ± SEM of Dynein binding, normalized to resting GST levels. **e**, Colocalisation between Bin1 and Dynein in cells stimulated with extra 10% serum (top), and between endophilin and dynein upon Amphiphysin/Bin1 double knock-down (DKD)(bottom). Right, histograms show the mean ± SEM from three independent biological experiments (*n*>50 puncta per condition). **f**, Model: Multi-layered regulation of FEME by Cdk5 and GSK3β: 1) obstruction of CRMP4 binding to Endophilin and thus PlexinA1 sorting into FEME carriers upon Semaphorin 3A stimulation, 2) inhibiton of Dynamin recruitment onto FEME carriers, thus inhibiting vesicle budding and 3) hinderance of Dynein recruitment by Bin1, thereby reducing FEME carriers movement. GSK3β binds to Endophilin and acts locally to hold off FEME. In cells exposed to growth factors, PI3K-mediated signaling activates AKT and other kinases that controls GSK3β activity, and thus license cells for FEME.

Pull-down experiments revealed that the SH3 domain of Bin1 and not that of Endophilin isolated endogenous Dynein (**Figure 7d**). As it was unlikely for Bin1 to bind directly to the motor domain, we tested Dynactin subunits that where identified by mass spectrometry in previous pull-downs^6^. However, neither Bin1 nor Endophilin bound to the p150^glued^ or p27 subunits (**Supplementary Figure S6e**). Stimulation of FEME by serum addition increased Dynein binding to Bin1 and recruitment onto FEME carriers (**Figure 7d-e**). Acute inhibition of Cdk5 and GSK3β increased the recruitment further, confirming that these kinases regulate Dynein loading onto FEME carriers (**Figure 6b, d and 7e**). Remaining EPAs produced in cells depleted for Bin1 had reduced levels of Dynein on them (**Figure 7e**), and often clustered at the cell periphery. Collectively, it showed that Bin1 recruits Dynein onto FEME carriers, under the control of Cdk5 and GSK3β. All our data taken together established the kinases as master regulators of FEME, antagonizing the process at several levels including cargo sorting, Dynamin and Dynein recruitment (**Figure 7f**).

## Discussion

Cdk5 and GSK3β play important roles in regulating endocytosis. The best understood mechanism involves the phosphorylation of Dynamin-1 at Ser778 by Cdk5, followed by that of Ser774 by GSK3β^9,18^. Phosphorylation of Ser778 hampers the recruitment of Dynamin-1 by binding partners such as Endophilin^20^. Phosphorylation of Ser774 inhibits Dynamin-1 activity^8^. Interestingly, these phosphorylations dampen CME but activate activity-dependent bulk endocytosis, suggesting they might mediate the crosstalk between the pathways^8,9^. Several other endocytic proteins, including Amphiphysin, are targets of the kinases in synapses^7,44^. The acute dephosphorylation of the formers upon axon depolarization is mediated by the Calcineurin phosphatase, which is activated by the sudden Ca^2+^ rise. Prompt removal of the inhibitory phosphorylations then swiftly activates compensatory endocytosis^45,46^.

Here, we found that, similarly to their function of regulating compensatory endocytosis in synapses, Cdk5 and GSK3β hold off FEME in non-neuronal cells (**Figures 1 to 4**). Some of the mechanisms are shared, as for example the regulation of Dynamin-1, but others appear different in the case of FEME. Indeed, the kinases also regulate cargo protein sorting by Endophilin, as well as Dynein recruitment by Bin1 (**Figures 5 to 7**). We established that Endophilin binds to PlexinA1 adaptor CRMP4 on a Proline-rich motif adjacent to the phosphorylation sites by Cdk5 and GSK3β, thereby placing the sorting of the receptor under negative regulation by the kinases (**Figure 5**). We also found the uptake of Slit1 and its receptor ROBO1 to be under control of the kinases for their recruitment into FEME carriers (**Supplementary Figure S4**). However, we do not know yet whether other FEME cargo generally adopts this regulation mechanism. Interestingly, there are several Serine-Proline motifs (consensus site for Cdk5) within or adjacent to the binding sites of Endophilin to several known FEME cargoes, including β1-Adrenergic receptor and CIN85 (the adaptor for EGFR).

In addition, the two kinases also block the recruitment of Dynein onto FEME carriers (**Figure 6**). We established that Bin1 engages the microtubule motor, and is another cytosolic marker of FEME carriers (**Figure 7**). Thus, like Endophilin, Bin1 functions in FEME in addition to its role in CME. Ubiquitously expressed short isoforms 9 and 10 colocalized poorly with CME markers, in agreement with their lack of binding motifs to AP2 and Clathrin^43^. We cannot rule out that Bin1 has additional functions other than recruiting Dynein, nor can we exclude that another protein recruits the motor protein onto EPAs either in the absence of, or in parallel to Bin1. How Cdk5 and GSK3β control the loading of FEME carriers onto Dynein is not known at this point. We tried obvious candidates p150^glued^ and p27, but they did not bind to Bin1. However, multiple Dynein adaptors such as Ndel1L, Lis1 or BICD, have been reported to be phosphorylated by either kinases^47,48,32^ and could be the link to Bin1. Additional layers of regulation are certainly at play as both Cdk5 and GSK3β regulate Dynein processivity, in addition to cargo loading^32,33,49,50^.

There may be additional levels of regulation by Cdk5 and GSK3β, either by regulation of other key steps of FEME or, more indirectly, by modulating the activity of other kinases. Interestingly, GSK3β inactivates both FAK1 (upon phosphorylation of Ser722 ^51^) and mTOR signaling (upon phosphorylation of TSC2 on Ser1341, Ser1337 and Ser1345 ^52^). Both FAK1 and mTORC1/2 were found to be FEME activators in our screen, as their acute inhibition blocked FEME (**Figure 1**).

The finding that Endophilin binds to GSK3β suggest that local regulation is required to control FEME. The binding to GSK3β but not α stems from the presence of several PRMs in the former that are not conserved in GSK3α. Even though GSK3α and β share high sequence similarity, no clear role for GSK3α has been assigned in endocytosis, perhaps owing to a defect in local recruitment, due to its lack of binding to Endophilin.

It is now clear that various cell types have different levels of FEME activity. There are strong differences in the maximum number in FEME carriers produced upon growth factor addition: a factor of 7 between the weakest and the strongest tested (HEK293 and HUVEC, respectively, **Supplementary Figure S1**). We also found that some cell types displayed spontaneous FEME (*i.e*. not induced experimentally) in some cell types, detected here in RPE1, hDFA and HUVEC cells. This is likely due to the high levels of growth factors in their culture in media: primary cells are routinely grown in medium supplemented with high doses of EGF, FGF, VEGF and/or IGF-1, which all trigger FEME^5^. Both Cdk5 and GSK3β control the level of FEME in a particular cell type, but only the sum of the activities of the two kinases (and perhaps that of other priming kinases that are yet to be identified) can predict the propensity of a cell type to be FEME active.

The signal relieving of the inhibition imposed by Cdk5 and GSK3β is not yet understood. The promptness of FEME activation upon stimulation of cargo receptors suggests that phosphatases likely erase inhibitory phosphorylations. However, which one could be acting downstream of receptors as diverse as Gα_s_ or Gα_i_-coupled GPCRs, RTKs, cytokine or cell guidance receptors (all the FEME cargoes known to date), is not obvious. In addition, kinases other than Cdk5 are likely priming GSK3β phosphorylation during FEME. Interestingly, some Endophilin functions are regulated by LRKK2, DYRK1A and Src^53-55^, but inhibitors toward these kinases did not affect spontaneous FEME in resting RPE1 cells. It is possible that dual inhibition together with GSK3β is required or that they regulate other FEME cargoes or processes. Thus, while a complex regulatory mechanism is likely to emerge from future work, the current study revealed the key role of Cdk5 and GSK3β in dampening FEME in absence of receptor activation.

## Supporting information

Ferreira et al. Supplementary Figures

## Author contributions

A.P.A.F., A.C., S.C.R., J.P., K.S. and E.B. performed biochemical assays; A.P.A.F., A.C., S.C.R., E.F.H. and E.B. performed cell biology experiments; J.T.K. provided guidance for some cell biology assays; S.S. performed mass spectrometry, under the supervision of K.T.; D.M. designed and performed FEME carrier isolation; A.P.A.F. and L.C.W.H. generated critical reagents; A.P.A.F., A.C., S.C.R., K.McG. and E.B. performed image acquisition and analysis; E.B. designed the research and supervised the project. E.B. wrote the manuscript with input from all the other authors.

## Acknowledgements

We thank Mina Edwards and Marta Martins for technical help, Alexandra Chittka (University College London), Tom Nightingale (Queen Mary University), Harvey McMahon (MRC Cambridge), Serge Benichou (Institut Cochin, Paris) and Michael Way (Crick Institute, London) for the kind gift of reagents and the members of the Boucrot lab for helpful comments. A.P.A.F was supported by the Fundação para a Ciência e Tecnologia. A.C. was supported by a Biotechnology and Biological Sciences Research Council (BBSRC) LIDo PhD scholarship. S.S. was supported by a Medical Research Council PhD scholarship. K.S. was a recipient of summer internship from the Lister Institute of Preventive Medicine. E.F.H. was the recipient of a Marie Skłodowska-Curie grant (661733). J.T.K. received support from an ERC starting grant (282430; Fuelling synapses) and from the MRC (MR/N025644/1). The G2-Si ion mobility mass spectrometer was purchased with a grant from the Wellcome Trust (104913/Z/14/ZBM) to K.T; and E.B. was a BBSRC David Phillips Research Fellow (BB/R01551X), a Lister Institute Research Fellow and a recipient of a BBSRC Pathfinder grant (BB/R01552X).

## Methods

### Cell culture

Human normal diploid hTERT-RPE-1 (ATCC CRL-4000, called ‘RPE1’ in this study) cells were cultured in DMEM:F12 HAM 1:1v/v (Sigma D6421), 0.25% Sodium bicarbonate w/v (Sigma), 1mM GlutaMAX-I (Thermo Fisher), 1X antibiotic-antimycotic (Thermo Fisher), and 10% Fetal Bovine Serum (FBS; Thermo Fisher). Human Primary Dermal Fibroblasts (ATCC PCS-201-012, called ‘hDFA’ in this study) were cultured in DMEM:F12 HAM 1:1v/v (Sigma D6421), 7.5mM GlutaMAX-I (Thermo Fisher), 1X antibiotic-antimycotic (Thermo Fisher), 2% Fetal Bovine Serum (FBS; Thermo Fisher), 0.8μM Insulin (MP Biomedicals 0219390025), 10ng/mL Basic Fibroblast Growth Factor (bFGF, LifeTech PHG0024), 50μg/mL Hydrocortisone and 1μg/mL ascorbic acid. Human umbilical vein endothelial cells, HUVEC (ATCC PCS-100-010 or a kind gift from Tom Nightingale (Queen Mary University)) were grown in endothelial cell growth medium containing 0.02mL/mL Fetal Calf Serum, 5ng/mL recombinant human Epidermal Growth Factor (EGF), 10ng/mL Basic Fibroblast Growth Factor (bFGF), 20ng/mL Insulin-like Growth Factor (IGF-1), 0.5ng/mL recombinant human Vascular Endothelial Growth Factor 165 (VEGF-165), 1μg/mL ascorbic acid, 22.5μg/mL Heparin and 0.2μg/mL Hydrocortisone (Promocell, C-22011). Human HeLa cells (a kind gift from Harvey McMahon), human embryonic kidney HEK293 (ATCC CRL-1573, called ‘HEK’ in this study) and African green monkey BSC-1 (ECACC 85011422), were cultured in DMEM (Sigma D6546) supplemented with 10% FBS, 1mM GlutaMAX-I (Thermo Fisher), 5% 1X antibiotic-antimycotic (Thermo Fisher). All the cells were maintained at 37°C, 5% CO2. Cells were regularly tested for mycoplasma contamination.

### *E.coli* BL21 (DE3)

For protein expression, *E.coli* BL21 (DE3) cells were grown in LB medium at 37°C.

### Small compound inhibitors and ligands

The following small compound inhibitors (amongst the best-reported inhibitors for each kinase ^11,12^) were used: AZ191 (called ‘DYRKi’ in this study Cayman 17693), AZD0530 aka Sacratinib (called ‘SRCi’ in this study, Cayman 11497), BI-D1870 (called ‘p90RSKi’ in this study, Cayman 15264), BIO-6-bromoindirubin-3′-oxime, aka BIO (called ‘GSK3i2’ in this study, (Sigma B1686), BI 2536 (called ‘PLKi’ in this study, Selleckchem S1109), CDK1/2 inhibitor III (called ‘Cdk1/2i’ in this study, Merck 217714), CHIR-99041 (called ‘GSK3i1’ in this study, Cayman 13122), Ciliobrevin D (called ‘Ciliobrevin’ in this study, Calbiochem 250401), CX-4945 (called ‘CK2i’ in this study, Cayman 16779), Dinaciclib (called ‘Cdk1/2/5/9i’ in this study, MedChemExpress Hy-10492), Dobutamine (Sigma D0676), GDC-0879 (called ‘BRAFi’ in this study, Tocris 4453), GDC-0941 (called ‘PI3Ki’ in this study, Symansis SYG0941), Genistein (called ‘Y-kinases’ in this study, Calbiochem 245834), GNE-7915 (called ‘LRRK2i’ in this study, MedChemExpress Hy-10328), GSK2334470 (called ‘PDKi’ in this study, Cayman 18095), GW 5074 (called ‘CRAFi’ in this study, Santa Crux sc-200639), JNK-IN-8 (called ‘JNKi’ in this study MedChemExpress Hy-13319), KT 5720 (called ‘PKAi’ in this study Cayman 10011011), MK2206 (called ‘AKTi’ in this study, LKT Laboratories M4000), MLR 1023 (called ‘LYNa’ in this study, Tocris 4582), PD0325901 (called ‘MEKi’ in this study, Tocris 4192), PD0332991 aka Palbociclib (called ‘Cdk4/6i’ in this study, Sigma PZ0199), PF-4708671 (called ‘p70S6Ki’ in this study, MedChemExpress Hy-15773), PF-4800567 (called ‘CK1Ei’ in this study, Cayman 19171), PHA-793887 (called ‘Cdk2/5/7i’ in this study, ApexBio A5459), PND-1186 (called ‘FAKi’ in this study, MedChemExpress Hy-13917), Purvalanol A (called ‘Cdk1/2/4i’ in this study, Santa Cruz sc-224244), P505-15 (called ‘SYKi’ in this study, Adooq Bioscence A11952), Roscovitine (called ‘Cdk1/2/5i in this study, Santa Cruz sc-24002), RO-3306 (called ‘Cdk1i’ in this study, Cayman 15149), SCH772984 (called ‘ERKi’ in this study, Sellekchem S7101), Staurosporine (called ‘broad kinases’ in this study, Alomone Labs AM-2282), STO609 (called ‘CaMKK1/2ii’ in this study, Cayman 15325), TAK-632 (called ‘panRAFi’ in this study, Selleckchem S7291), Torin 1 (called ‘mTORC1i’ in this study, Tocris 4247), VX-745 (called ‘p38i’ in this study, MedChemExpress Hy-10328) and ZM 447439 (called ‘AurA/AurBi’ in this study, Cayman 13601). The following ligands were used: human Semaphorin 3A extracellular region 6 (6xN-terminal His-tag, R&D 1250-S3) and human Slit1 (6xC-terminal His-tag, R&D 6514-SL-050).

### Gene cloning and mutagenesis

Full length and truncated genes (all human, unless specified) were amplified and cloned into pDONR201 (Invitrogen) and transferred into pEGFP, pTagRFP-T (called ‘RFP’ elsewhere), pMyc or pGEX-6P2 vectors converted into the Gateway system (pDEST vectors made from a pCI backbone), as appropriate: Endophilin-A2 (*SH3GL1*, IMAGE 3458016) full length and SH3 domain (aa 311-end); Endophilin-A1 (*SH3GL2 iso1*, FLJ 92732) full length and SH3 domain (aa 295-end); Endophilin-A3 (*SH3GL3* iso 1, IMAGE 5197246) full length and SH3 domain (aa 291-end); Bin1, also known as Amphiphysin-II (*BIN1 iso9*, cloned from human brain cDNA library) full length; full length CRMP2 (*DPYSL2*, DNASU HsCD00513405), full length CRMP3 (*DPYSL4*, NM_006426 Origene), full length mouse CRMP4 (*DPYSL3*, Origene 1197294), full length CRMP5 (*DPYSL5*, amplified from human brain cDNA library, Novagen), Ephrin receptor A1 cytoplasmic tail (aa 568-976) (*EPHA1*, DNASU HsCD00516390), Ephrin receptor A6 cytoplasmic tail (aa 572-1036) (*EPHA6*, DNASU HsCD00350501), Ephrin receptor B1 cytoplasmic tail (aa 259-346) (*EPHB1*, DNASU HsCD00038738), Ephrin receptor B4 cytoplasmic tail (aa 561-987) (*EFNB4*, DNASU HsCD00021508), Ephrin receptor B6 cytoplasmic tail (aa 616-1021) (*EPHB6*, DNASU HsCD00505529), Semaphorin 4F cytoplasmic tail (aa 681-770) (*SEMA4F*, DNASU HsCD00041427); Semaphorin 6A cytoplasmic tail (aa 671-1030) (*SEMA6A*, Sino Biologica HG11189-M); Semaphorin 6B cytoplasmic tail (aa 616-1021) (*SEMA6B*, amplified from human brain cDNA library, Novagen); Semaphorin 6D cytoplasmic tail (aa 684-1073) (*SEMA6D*, DNASU HsCD00516397); Plexin B1 cytoplasmic tail (aa 1512-2135) (*PLXNB1*, Addgene 25252), mouse Roundabout homolog 1 cytoplasmic tail (aa 880-1612) (*ROBO1*, DNASU HsCD00295416); Roundabout homolog 3 cytoplasmic tail (aa 912-1386) (*ROBO3*, DNASU HsCD00302878) and Netrin receptor UNC5B cytoplasmic tail (aa 398-945) (*UNC5B*, DNASU HsCD294959), EGFP-p27 (Addgene #15192); EGFP-p150Glued (Addgene #36154). Bovine Dynamin 1-EGFP and rat GST-Bin1 SH3 domain were kind gifts from Harvey McMahon (MRC Cambridge), EGFP-p50 dynamitin (full length *DCTN2*) was a kind gift from Serge Benichou (Institut Cochin, Paris) and EGFP-TPR (mouse KLC2 TPR domains aa 155-599) was a kind gift from Michael Way (Crick Institute, London). EGFP-tagged human full-length BAR domain proteins library was described before^6^. Point mutations P502A, S522D, S522A, R525E and P526A were introduced in full length CRMP4 and S774A and S778A were introduced in full length Dynamin-1 by site-directed mutagenesis and verified by sequencing.

### Gene transfection

For fixed cell colocalization experiments, cells seeded on 13mm coverslips (placed in 24-well plates) were transfected using Lipofectamine 2000 (Thermo Fisher) or Nanofectin (PAA) and 10 to 500ng DNA depending on the plasmids and the experiments (low or high overexpression). The levels of each plasmid were titrated down to low levels allowing good detection but limiting side effects of overexpression. Cells seeded onto live-cell imaging 35 mm glass bottom dishes (MatTek) were transfected using Lipofectamine 2000 (Thermo Fisher) and 50 to 250ng DNA. For pull-down experiments, co-immunoprecipitation and EGFP-trap immunopurifications, HEK293 cells seeded in 6-well plates or 100mm dishes were transfected using GeneJuice (Merck) and 1 to 3µg DNA. Cells were incubated 16 to 24h to express the constructs and were either imaged live, fixed (4% pre-warmed paraformaldehyde, 20min at 37°C) or processed to prepare cell extracts.

### siRNA suppression of gene expression

The following siRNA oligos (all Stealth, Thermo Fisher) were used: Endophilin A1, A2 and A3 triple knock-down (Endo TKD) was achieved by combining oligos against Endophilin A1 (Thermo HSS109709; 2 oligos against human *SH3GL2*), Endophilin A2 (Thermo HSS109707; 2 oligos against *SH3GL1*) and Endophilin A3 (Thermo HSS109712; 2 oligos against human *SH3GL3*); AP2: HSS101955 (2 oligos against human *AP2M1*); CDK5 (Thermo HSS101729; 2 oligos against human *CDK5*); GSK3α/β double knock-down (DKD) was achieved by combining oligos against GSK3α (Thermo HSS104518; 2 oligos against human *GSK3A*) and GSK3β (Thermo HSS104522; 2 oligos against human *GSK3B*); CDK5+GSK3α/β triple knock-down (TKD) was achieved by combining aforementioned oligos against CDK5 and GSK3α*/*β; AMPH+Bin1 double knock-down (DKD) was achieved by combining oligos against Amphiphysin-1 (Thermo HSS100465; 2 oligos against human *AMPH*) and Bin1 (Thermo HSS100468; 2 oligos against human *BIN1*); FBP17+CIP4+TOCA-1 triple knock-down (TKD) was achieved by combining oligos against <u>FBP17</u> (Thermo HSS118093; 2 oligos against human *FNBP1*), <u>CIP4</u> (Thermo HSS113814; 2 oligos against human *TRIP10*) and <u>TOCA-1</u> (Thermo HSS123422; 2 oligos against human *FNBP1L*) and <u>Lamellipodin</u>: Dharmacon ON-TARGETplus SMARTpool (mix of J-031919-08, J-031919-07, J-031919-06 and J-031919-05 targeting human *RAPH1*). Control siRNA used were Invitrogen Stealth control (scrambled) oligo 138782. Cells seeded on 13 mm coverslips placed in 24 well plates were transfected twice (on day 1 and 2) with Oligofectamine or RNAi MAX (Thermo Fisher) complexed with 20pmol of each indicated siRNA and analyzed 3 to 4 days after the first transfection. RNAi knock-down efficiency was verified by western-blotting or immunofluorescence counter-staining. The use of validated pools of siRNA targeting the same genes increased the knock-down efficiency and specificity.

### Antibodies

The following antibodies were used for immunostaining or immunoblotting: anti-EGFP ab290 (rabbit polyclonal, AbCam290), anti-EGFP clones 7.1 and 13.1 (mouse monoclonal, Roche 11814460001), anti-Endophilin A2 clone H-60 (rabbit polyclonal, Santa Cruz 25495), anti-Endophilin A2 clone A-11 (mouse polyclonal, Santa Cruz 365704), anti-β1 adrenergic receptor (rabbit polyclonal, AbCam ab3442), anti-CRMP4 (rabbit polyclonal, Milipore 5454), anti-Dynein clone 74.1 (mouse monoclonal, eBioscience 14-9772-80), anti-Plexin A1 (rabbit polyclonal recognizing the ectodomain of PlexinA1, Alome labs Ab32960), anti-ROBO1 (sheep polyclonal, AF7118 R&D Systems), anti-LAMP-1 (mouse monoclonal clone H4A3-c, Developmental Studies Hybridoma Bank), anti-phosphorylated Ser9 GSK3β clone D85E12 (rabbit monoclonal, Cell Signaling Technology 5558), anti-GSKα*/*β D75D3 (rabbit polyclonal, Cell Signaling Technology 5676), anti-Dynamin 1 clone 41 (mouse monoclonal, BD Pharmigen 610245), anti-Bin1 (rabbit polyclonal, GeneTex GTX103259), anti-Lamellipodin (rabbit polyclonal, Atlas Antibodies HPA020027), anti-CIP4, (mouse monoclonal clone 21, Santa Cruz sc-135868), anti-α Tubulin clone TUB2.1 (mouse monoclonal, AbCam ab11308) and anti-His tag clone D3I10 (rabbit polyclonal, Cell Signaling Technology 12698). The following secondary antibodies were used for microscopy: Alexa Fluor 488 and 555 goat anti-mouse IgG, Alexa Fluor 488 and 555 goat anti-rabbit IgG, Alexa Fluor 388 Donkey anti-Sheep IgG and Fluor 555 donkey anti-mouse IgG (all from Life technologies). For immunoblot; goat anti-mouse IgG-HRP conjugate and goat anti-rabbit IgG-HRP conjugated (both from Bio-Rad). Actin was stained using Phalloidin-Alexa647 (Cell Signaling Technology 8940) and DNA using DRAQ5 (BioStatus DR50200).

### Cell stimulation and cargo uptake

Cells were kept at 37°C and 5% CO_2_ during the whole assay (apart during medium exchanges) and never serum-starved or pre-incubated at 4°C. ‘Resting’ conditions correspond to cells being cultured in 10% serum media and directly fixed (4% pre-warmed paraformaldehyde) for 20min at 37°C. Kinase inhibition was achieved by incubating cells grow in full medium (10% serum) with the indicated small compound inhibitors at the indicated concentrations and for the indicated times at 37°C before being washed once with pre-warmed PBS and fixed (4% pre-warmed paraformaldehyde) for 20min at 37°C. Serum stimulation was achieved by adding 37°C pre-warmed 10% serum on complete medium (20% serum final) for the indicated times. β1 adrenergic receptor stimulation (which activates FEME) was performed by incubating cells at 37°C for 4 or 30min with pre-warmed medium containing 10μM dobutamine. Plexin A1 uptake was performed by incubating cells at 37°C for 5 to 20min with pre-warmed medium containing 20nM Semaphorin 3A and 10 μg/mL anti-PlexinA1 antibodies (recognizing the ectodomain of PlexinA1). ROBO1 stimulation was performed by incubating cells at 37°C for 10min with pre-warmed medium containing 2nM Slit1-(His)_6_. In some experiments, cells were pre-incubated at 37°C for the indicated times with small compound inhibitors before stimulation with dobutamine, Semaphorin 3A or Slit1 (in constant inhibitor concentration). After the incubation periods at 37°C, cells stimulated as described above were quickly washed once with 37°C pre-warmed PBS to removed unbound ligands and fixed with pre-warmed 4% PFA for 20min at 37°C (to preserve Endophilin staining and FEME carriers morphology). In some experiemnts unbound and cell suface anti-PlexinA1 antibodies were removed by one quick wash in ice-cold PBS^++^ (containing 1mM CaCl_2_ and 1mM MgCl_2_) followed by two 5 min incubations in acid stripping buffer (150mM NaCl, 5mM KCl, 1mM CaCl_2_, 1mM MgCl_2_, 0.2M acetic acid adjusted to pH2.5), followed by two wash in PBS^++^ to normalize pH back to 7. Fixed cells were then washed three times with PBS and one time with PBS supplemented with 50mM NH_4_Cl to quench free PFA. Cells were then permeabilized (0.05% saponin), immunostained and imaged as described below.

### Immunostaining and confocal fluorescence microscopy

Cells were fixed with 4% PFA at 37°C for 20 minutes, washed 3 times with PBS and 1 time with PBS with 50 mM NH_4_Cl to quench free PFA. Cells were then permeabilized for 5 minutes with PBS with 0.05% saponin and immunostained with primary and secondary antibodies in PBS with 0.05% saponin (Sigma) and 5% heat inactivated Horse Serum. Cover slips 0.13-0.16 mm (Academy) were mounted on slides (Thermo scientific) using immunomount DAPCO (GeneTex) and imaged using a laser scanning confocal microscope (TCS Sp5 AOBS; Leica) equipped with a 63x objective. For Alexa488, the illumination was at 488nm and emission collected between 498 and 548nm; for Alexa555 the laser illumination was at 543nm and emission collected between 555 and 620nm; for Alexa647 and DRAQ5, the laser illumination was at 633nm and emission collected between 660 and 746nm. Correlation between total Cdk5 or GSK3β cellular levels and number of EPAs were determined from single cell measurements the same cells (*i.e*. matching Cdk5 or GSK3β levels with the number of EPAs in each individual cell measured). The percentages of endophilin spots located at the leading edge of cells or FEME carriers (EPAs) positive for endogenous GSK3β, Dynamin-1, CRMP4, Dynein or Bin1 were determined by line scans using Volocity 6.0. as previously described^5,6^. Colocalization of overexpressed Endophilin-A2-RFP and EGFP-tagged CRMP4 constructs were determined by Manders’ overlap using Volocity 6.0. The percentages of FEME carriers (EPAs) positive for BAR domain tagged with EGFP were determined by line scans using Volocity 6.0. as previously described^5,6^. The percentages of Bin1 spots positive for endogenous Endophilin, Lamellipodin, CIP4 or Dynein were determined by line scans using Volocity 6.0. as previously described^5,6^.Levels of endogenous β1AR, PlexinA1 or recombinant Slit1 internalized into FEME carriers were measured by using Volocity 6.0. as previously described^5,6^.

### Live-cell confocal fluorescent microscopy

Just before live-cell imaging, the medium of cells grown on MatTek dishes was changed to α-MEM without phenol red, supplemented with 20mM HEPES, pH7.4 and 5% FBS and placed into a temperature controlled chamber on the microscope stage with 95% air: 5% CO_2_ and 100% humidity. Live-cell imaging data were acquired using a fully motorized inverted microscope (Eclipse TE-2000, Nikon) equipped with a CSU-X1 spinning disk confocal head (UltraVIEW VoX, Perkin-Elmer, England) using a 60x lens (Plan Apochromat VC, 1.4 NA, Nikon) under control of Volocity 6.0 (Improvision, England). 14-bit digital images were obtained with a cooled EMCCD camera (9100-02, Hamamatsu, Japan). Four 50mW solid-state lasers (405, 488, 561 and 647nm; Crystal Laser and Melles Griots) coupled to individual acoustic-optical tunable filter (AOTF) were used as light source to excite EGFP and TagRFP-T. Rapid two-colour time-lapses were acquired at 500ms to 2s intervals, using a dual (525/50; 640/120, Chroma) emission filter respectively. The power of the lasers supported excitation times of 50ms in each wavelength and the AOTFs allowed minimum delay (∼1ms) between 2 colors (e.g. delay between green-red for each timepoint), which was an important factor to assess the colocalization between markers.

### Protein purification and pull down experiments

GST or GST-tagged SH3 domains were expressed in BL21 (DE3) *E.coli* (New England Biolabs). Cells were lysed by sonication in presence of lysozyme (Affymetrix), protease inhibitor (Thermo Scientific) and DNAse powder (Sigma-Aldrich), spun at 11,000*g* for 1h at 4°C. The supernatants containing the GST or GST-SH3 domains (soluble fraction) were concentrated (to ∼ 60 mg/mL) using a Centricon Plus 70-1000 NMWL (Centricon) for 1h at 4°C and then incubated rotating with GST-sepharose beads (PierceTM glutathione superflow agarose) overnight at 4°C. The beads were washed 10 times with ice-cold PBS and kept in PBS and sodium azide 0.02% solution at 4°C and used in pull-down assays. Cell lysates - non-transfected or overexpressing EGFP-tagged proteins -were prepared in lysis buffer (20mM HEPES, 1mM EDTA, 0.2% Triton X-100 and protease inhibitor cocktail (Roche)) briefly sonicated (three times 5 second pulses with 30 seconds rest, 10 μm amplitude) and spun at 20,000 *g* for 10 min at 4°C. Cell lysates were incubated with bead-bound proteins (amounts were qualibrated by gel electrophoresis followed by Coomasie to equivalent amounts) overnight at 4°C and then centrifuged at 7,500*g* and washed 3 times with lysis. The remaining bead pellet was boiled in sample buffer and run on SDS-PAGE. Input lanes correspond to 5% of cell extract. The final pellets (bound fractions), supernatants (unbound fractions) and original extracts (input fractions) were boiled in sample buffer and ran on SDS-PAGE. The proteins were transferred onto PVDF membrane and immunoblotted using anti-EGFP antibodies or antibodies against endogenous proteins, as indicated, followed by HRP-coupled secondary antibodies (BioRad). Blots were developed with the ECL kit (Thermo Fischer Scientific or Merck Millipore) and x-ray film and quantified using ImageLab.

### Co-immunoprecipitations

HEK293 cells were co-transfected with equal amounts (1 to 3μg) of Myc- and EGFP-tagged BAR domain constructs. After 16-24h expression, cells were quickly washed with cold PBS, lysed in ice-cold lysis buffer (10mM Tris HCL pH7.5, 150mM NaCl, 0.5mM EDTA, 0.5% NP40 and a protease and phosphatase inhibitor cocktail (Thermo Scientific)) and spun at 14,000*g* for 10min at 4°C. Cell lysates were incubated with GFP-TRAP_A or M (Chromotek) bead slurry for 1 to 16h at 4°C. The beads were washed 3 times (10mM Tris HCL pH7.5, 150mM NaCl, 0.5mM EDTA). The final pellets and unbound fractions were boiled in SDS sample buffer and ran on SDS-PAGE (‘input’ lanes correspond to 1 to 10% of cell extracts). The proteins were transferred onto PVDF membrane and immunoblotted using anti-Myc, anti-EGFP, antibodies or antibodies against endogenous proteins, as indicated, followed by HRP-coupled secondary antibodies (BioRad). Blots were developed with the ECL kit (Thermo Fischer Scientific or Merck Millipore) and x-ray films.

### FEME carrier isolation

RPE1 cells grown on 15 cm dishes were stimulated with extra 10% FBS (20% final) for 10 min at 37°C, quickly rinsed with ice-cold PBS^+++^ (PBS with 1mM Ca2+, protease and phosphatase inhhibitors), collected using a cell scraper and pelleted (400*g*, 5min, 4°C). Cell pellets were loosened in 1mL of ice-cold homogenization buffer A (HBA) (3 mM Imidazol pH 7.4, 1 mM EDTA, and 0.03 mM cycloheximide plus protease and phosphatase inhibitor cocktail), spun (1,300*g*, 10min, 4°C), resuspendend into one volume of HBA and incubated for 20min on ice. One volume of homogenization buffer B (HBB; HBA containing 500mM sucrose) was added and cells were mechanically lysed through 25G needles, avoiding nuclei disruption. Homogenates were diluted into HBB (one part homogenetate and 0.7 part HBB) and post-nuclear supernatants (PNS) were collected after spinning (two time, 2,000*g*, 10min, 4°C). Sucrose concentration of the PNS was adjusted to 40.6% using 62% sucrose solution (2.351 M sucrose, 3 mM Imidazole pH 7.4) and 1 volume was loaded at the bottom of ultracentrifuge tubes. Cushions of 35% sucrose (1.5 volume), 25% (1 volume) and 8% sucrose (1 volume) were carefully added and the tubes were centrifuged at 210,000*g* for 3h at 4°C. Gradients were divided in 10 fractions and used either for immunoblotting of immunopurification. Immunobloting was performed after sucrose gradient fractions were concentrated following protein precipitation (25%/v TCA, incubated for 10min at 4°C, spun at 20,000*g* for 5min at 4°C; pellets were washed twice with -20°C pre-chilled acetone and air dried before resuspension in SDS sample buffer). Selected sucrose gradient fractions were submitted to immunoprecipitation using an anti-Endophilin antibody (mouse IgG2a/?-ligh chain clone A-11, sc365704) coupled directly to hydrazide-terminated magnetic beads (Bioclone Inc; coupling performed following manufacturer’s instructions) to allow for elution of the binding material without denaturation. Selected sucrose gradient fractions were adjusted to 500 µl with HBA and incubated (overnight at 4°C, slow rotation) with 1:50^th^ volume of anti-Endophilin coupled magnetic beads (pre-equilibrated in HBA with 250 mM sucrose). Unbound material was isolated, and the beads washed three times in 1mL ice-cold HBB (washes were also kept for analysis). Bound material was released in 500mL elution buffer (0.1 M glycine pH 2, 250 mM sucrose) for 10 minutes (4°C, slow rotation), neutralized (>200µl of 1 M Tris pH 8 until pH back to neutral) and prepared for immunoblotting. Lipids were stained using alcohol-free coomassie, as established previously^56^.

### Immunoprecipitation for MS experiments

RPE1 cells were grown on 10cm dishes to a confluence of 80% before harvest. For Stimulated condition, cell media was supplemented with extra 10% FBS (20% final) for 5 min at 37°C. For Resting condition, no additional treatment was performed prior to cell lysis. Cells were washed 2 times in ice-cold PBS before being gently scraped into lysis buffer (10 mM Tris HCL pH7.5, 150 mM NaCl, 0.5 mM EDTA, 0.5% NP40 and a protease and phosphatase inhibitor cocktail (Thermo Scientific)) and incubated on ice for 30 min, then centrifuged at 17,000 x g for 10 min. Anti-endophilin antibodies (Endophilin II A-11) were coupled in-house to hydrazide-terminated magnetic beads (H-beads). 10 µl of H-beads pre-washed in lysis buffer were incubated with 470 µl of cell lysate overnight at 4°C with end-over-end rotation. The beads were washed 3 times in lysis buffer then boiled in 50 µl SDS sample buffer for 10 min.

### Sample preparation by in-gel digestion

Samples were separated by SDS-PAGE on a 4-12% gel and stained with InstantBlue (Expedeon) staining solution. Sample lanes were cut into 10 sections, then further cut into 1mm^3^ pieces and washed in destaining solution (40% ethanol, 10% glacial acetic acid in water). Proteins were reduced in 10 mM DTT, then alkylated with 20 mM iodoacetamide in 50mM ammonium bicarbonate buffers. Gel pieces were washed then immersed in a 10 ng/µl Trypsin buffer in 50 mM ammonium bicarbonate and digested for 16-18 hours at 37°C. Gel pieces were incubated in Elution buffer (1% formic acid, 2% acetonitrile in LC-MS grade water (Thermo Scientific)) and dried in a SpeedVac. Peptides resuspended in 0.5% acetic acid in water and desalted on C18-Stagetips, then dried in a SpeedVac. Peptides were resuspended in LC-MS running buffer (3% acetonitrile, 0.1% Formic acid in water) prior to analysis by LC-MS and spiked with *E. coli* ClpB peptides (Waters, UK) such that 50 fmol of spiked-in peptide standard was introduced per injection.

### Liquid chromatography and mass spectrometry data acquisition

LC-MS grade solvents were used for all chromatographic steps. Separation of peptides was performed using a Waters NanoAcquity Ultra-Performance Liquid Chromatography system. Peptides were reconstituted in 97:3 H2O:acetonitrile + 0.1% formic acid. The mobile phase was: A) H2O + 0.1% formic acid and B) Acetonitrile + 0.1% formic acid. Desalting of the samples was performed online using a reversed-phase C18 trapping column (180 µm internal diameter, 20 mm length, 5 µm particle size; Waters). Peptides were separated by a linear gradient (0.3 µl/min, 35°C column temperature; 97-60% Buffer A over 60 minutes) using an Acquity UPLC M-Class Peptide BEH C18 column (130Å pore size, 75µm internal diameter, 250mm length, 1.7µm particle size, Waters, UK). [Glu1]-fibrinopeptide B (GFP, Waters, UK) was used as lockmass at 100 fmol/µl. Lockmass solution was delivered from an auxiliary pump operating at 0.5 µl/min to a reference sprayer sampled every 60 seconds. The nanoLC was coupled online through a nanoflow sprayer to a QToF hybrid mass spectrometer (Synapt G2-Si; Waters, UK). Accurate mass measurements were made using a data-independent mode of acquisition (HDMS^E^). Each sample was analysed in technical duplicate.

### Database searching

Raw data was analyzed using Progenesis v4.0 (Waters, UK). Data were queried against a Homo sapiens FASTA protein database (UniProt proteome:UP000005640) concatenated with a list of common contaminants obtained from the Global Proteome Machine (ftp://ftp.thegpm.org/fasta/cRAP) and *E. coli* ClpB, which acted as a standard for label-free absolute protein quantitation ^57^. Carbamidomethyl-C was specified as a fixed modification and Oxidation (M) and Phosphorylation of STY were specified as variable modifications. A maximum of 2 missed cleavages were tolerated in the analysis to account for incomplete digestion. For peptide identification 3 corresponding fragment ions were set as a minimum criterion whereas for protein identification a minimum of 7 fragment ions were required. Protein false discovery rate was set at 1%. Samples were normalized to Endophilin peptide abundance and fractions were combined *in-silico* in Progenesis to obtain absolute protein abundances for differential expression analysis.

### Experimental Design

A strategy for randomization, stratification or blind selection of samples has not been carried out. Sample sizes were not chosen based on pre-specified effect size. Instead, multiple independent experiments were carried out using several independent biological replicates as detailed in the figure legends.

### Quantification and Statistical Analysis

All experiments were repeated at least three times, giving similar results. For all figures, results shown are mean ± standard error of the mean (SEM). Statistical testing was performed using Prism 6 (GraphPad Software). Comparisons of data were performed by one-way analysis of variance (ANOVA) with Tukey’s multiple comparison test or by two way ANOVA with Tukey’s multiple comparisons test, as appropriated. *NS*, non significant (P>0.05); *, *P*<0.05, **, *P* <0.01, ***, *P* <0.001.

## Supplementary Figure Legends

**Supplementary Figure 1**. *Related to Figures 1 and 2*. **a**, Spontaneous FEME carrier formation in HeLa, HEK293, BSC1, RPE1, human primary dermal fibroblasts (hDFA) or HUVEC cells grown in their respective complete culture media (see Methods). Arrowheads point at FEME carriers. **b**, Histograms show the mean ± SEM.of the percentage of resting or stimulated (+10% FBS) cells displaying active FEME (n>50 cells per condition, from independent biological triplicate). **c**, Histograms show the mean ± SEM of the number of FEME carriers in resting or stimulated (+10% FBS) cells (n>150 EPAs per condition, from independent biological triplicate). Statistical analysis was performed by two-way ANOVA; *NS*, non significant; *, *P* <0.05, **, *P* <0.01, ***, *P* <0.001. Scale bar, 20μm.

**Supplementary Figure 2**. *Related to figure 3*. **a**, Volcano plot (-Log10 of *p* values versus Log2 of fold changes in proteins levels; Log2 of protein abundances are shown as heat map representation) of proteins detected by mass spectrometry from fractions immunoprecipitated with anti-Endophilin antibodies. Cells were stimulated with additional 10% serum (FBS) or not (resting) prior to extraction and co-immunoprecipitation. Proteins relevant to this study were annotated. Full list of the proteins detected in Supplementary Table 1. **b**, Confocal images showing the levels of Endophilin and Cdk5 or GSK3 a/β in cells depleted of Cdk5 (CDK5 KD) or GSK3α/β (GSK3α/β DKD), respectively. Arrowheads point at FEME carriers. Scale bars, 20μm.

**Supplementary Figure 3**. *Related to Figure 5*. **a**, Quantification of experiments illustrated in Figure 5b. Histograms show the mean ± SEM from three independent biological experiments. **b**, Quantification of experiments illustrated on Figure 5c. Histograms show the mean ± SEM from three independent biological experiments. **c**, Quantification of experiments illustrated on Figure 5d. Histograms show the mean ± SEM from three independent biological experiments. **d**, Recruitment of the indicated overexpressed EGFP-tagged CRMP4 constructs onto structures formed by overexpressed Endophilin-A2-RFP. Arrowheads point at overexpressed Endophilin-RFP structures. Histograms show the mean ± SEM from biological triplicates (*n*=15 cells per condition). **e**, Internalized Plexin A1 (whole uptake, FEME plus other pathways) in control cells or cells depleted for CRMP4, AP2 or endophilin A1, A2 and A3 (‘Endo TKD’), upon stimulation with 20nM Semaphorin 3A with 10μg/mL anti-PlexinA1 antibodies (recognizing the ectodomain of PlexinA1) for 20 minutes. Unbound and cell surface bound anti-PlexinA1 antibodies were removed prior to fixation. **f**, Related to Figure 5g: whole dataset (note that some images are similar, as Figure 5g shows a subset of the conditions tested). Endogenous Plexin A1 uptake into FEME carriers in HUVEC cells depleted of Endophilin A1, A2 and A3 (‘Endophilin TKD), Cdk5 and GSK3α and β (‘CDK5+GSK3α/β TKD’), CRMP4 or AP2 or pre-treated with Cdk5i and/or GSK3i for 5min. Cells were stimulated by 20nM Semaphorin 3A (Sema3A) for 5min in presence of 10 μg/mL anti-PlexinA1 antibodies (recognizing the ectodomain of PlexinA1) or not (resting). Arrowheads point at FEME carriers. Statistical analysis was performed by one-way ANOVA (a, b, d and e) or two-way ANOVA (c); *NS*, non significant; *, *P*<0.05, **, *P* <0.01, ***, *P* <0.001. Scale bars, 20 (d) and 5μm (f).

**Supplementary Figure 4**. *Related to figure 5*. **a**, Left, pull-down using GST-SH3 domains of Endophilin A1, A2 or A3 and cell extracts expressing the indicated EGFP-tagged receptor tailss. GST was used as negative control. Binding proteins were detected by immunoblotting with an anti-EGFP antibody. ‘input’ lanes correspond to 5% of the cell extracts. Right, Histograms show the mean ± SEM from three independent biological experiments. **b**, Slit1-His_6_ uptake (2nM for 5min) into FEME carriers in HUVEC cells depleted of Endophilin A1, A2 and A3 (‘Endophilin TKD), Cdk5 and GSK3α and β (‘CDK5+GSK3α/β TKD’), or AP2 or pre-treated with 5μM Dinaciclib (Cdk5i) and/or CHIR-99021 (GSK3i) for 10min. Arrowheads point at FEME carriers. Histograms show the mean ± SEM from biological triplicates (*n*=15 cells per condition). Statistical analysis was performed by one-way ANOVA (a) or two-way ANOVA (b); *NS*, non significant; *, *P*<0.05, **, *P* <0.01, ***, *P* <0.001. Scale bar, 5μm.

**Supplementary figure 5**. *Related to figure 6*. **a**, FEME carrier formation in RPE1 cells treated for 10min with 5μM Ciliobrevin (Dynein inhibitor) or overexpressing EGFP-tagged p50 dynamitin (Dynein dominant-negative) or Kinesin TPR domain (Kinesin dominant-negative), followed by 10μM dobutamine for 4min. EGFP was used as negative control. Arrowheads point at FEME carriers. Histograms show the mean ± SEM from biological triplicates (*n*=20 cells per condition). **b**, Lysosomal accumulation of β1 adrenergic receptor (β1AR) RPE1 cells overexpressing EGFP-tagged p50 dynamitin or Kinesin TPR domain and treated with 10μM dobutamine for 30min. Arrowheads point at β1AR inside lysosomes. Histograms show the mean ± SEM from biological triplicates (*n*=30 cells per condition), normalized to control cells.

**Supplementary figure 6**. *Related to figure 7*. **a**, Colocalization of endogenous Bin1 and AP-2 or Clathrin upon stimulation with 10μM dobutamine for 4min. Right, histograms show the mean ± SEM from three independent biological experiments (*n*>150 puncta per staining). Arrowheads point at Bin1 spots. b**e**, Colocalization of endogenous Bin1 and Lamellipodin in cells depleted of Lamellipodin (‘Lpd KD’), Bin1 Amphiphysin (‘Amph+Bin1 DKD’), or not (‘resting’). Arrowheads point at Bin1 or Lpd spots at the plasma membrane. Right, histograms show the mean ± SEM from three independent biological experiments (*n*>150 puncta per condition). **c**, Colocalization of endogenous Bin1 and CIP4 in cells depleted of FBP17, CIP4 and TOCA-1 (‘FCT TKD’), Bin1 Amphiphysin (‘Amph+Bin1 DKD’), or not (‘control’), and stimulated with 10μM dobutamine for 4min. Right, histograms show the mean ± SEM from three independent biological experiments (*n*>150 puncta per condition). **d**, Pull-down experiments using beads with GST-SH3 domains of Endophilin A2 or Bin1, in resting cells or cells treated with extra 10% FBS for 10 min. GST beads were used as negative control. Inputs correspond to 4% of cell extracts. Bottom, histograms show the mean ± SEM of Dynein binding, normalized to resting GST levels. **e**, Pull-down experiment using beads with GST only or GST-SH3 domains of endophilin A2 or Bin1. Bound EGFP, p150-glued-EGFP, p27-EGFP or Dynamin 1-EGFP were tested by immunoblot. GST was used as negative control. Input corresponds to 4% of cell extracts. Statistical analysis was performed by one-way ANOVA; *NS*, non significant; *, *P*<0.05, **, *P* <0.01, ***, *P* <0.001. Scale bars, 5 (a) and 20μm (b).

**Supplementary Table 1**. List of interactors identified in Supplementary Figure 2a. Proteins co-immunoprecipirating with Endophilin from resting or FBS-stimulated RPE1 cells (+10% additional FBS in regular media for 10min) were identified by mass spectrometry.

## References

1 McMahon, H. T. & Boucrot, E. Molecular mechanism and physiological functions of clathrin-mediated endocytosis. Nat Rev Mol Cell Biol 12, 517–533 (2011).

2 Kaksonen, M. & Roux, A. Mechanisms of clathrin-mediated endocytosis. Nat Rev Mol Cell Biol 19, 313–326 (2018).

3 Ferreira, A. P. A. & Boucrot, E. Mechanisms of carrier formation during clathrin-independent endocytosis. Trends in Cell Biology 28, 188–200 (2018).

4 Sandvig, K., Kavaliauskiene, S. & Skotland, T. Clathrin-independent endocytosis: an increasing degree of complexity. Histochem Cell Biol (2018).

5 Boucrot, E. et al. Endophilin marks and controls a clathrin-independent endocytic pathway. Nature 517, 460–465 (2015).

6 Chan Wah Hak, L. et al. FBP17 and CIP4 recruit SHIP2 and lamellipodin to prime the plasma membrane for fast endophilin-mediated endocytosis. Nat Cell Biol 20, 1023–1031 (2018).

7 Liang, S. et al. Major Cdk5-dependent phosphorylation sites of amphiphysin 1 are implicated in the regulation of the membrane binding and endocytosis. J Neurochem 102, 1466–1476 (2007).

8 Reis, C. R. et al. Crosstalk between Akt/GSK3beta signaling and dynamin-1 regulates clathrin-mediated endocytosis. EMBO J 34, 2132–2146 (2015).

9 Clayton, E. L. et al. Dynamin I phosphorylation by GSK3 controls activity-dependent bulk endocytosis of synaptic vesicles. Nat Neurosci 13, 845–851 (2010).

10 Smillie, K. J. & Cousin, M. A. Akt/PKB controls the activity-dependent bulk endocytosis of synaptic vesicles. Traffic 13, 1004–1011 (2012).

11 Wang, J. & Gray, N. S. SnapShot: Kinase Inhibitors I. Mol Cell 58, 708 e701 (2015).

12 Wang, J. & Gray, N. S. SnapShot: Kinase Inhibitors II. Mol Cell 58, 710 e711 (2015).

13 Malumbres, M. Cyclin-dependent kinases. Genome Biol 15, 122 (2014).

14 Cohen P F. S. The renaissance of GSK3. Nat Rev Mol Cell Biol 2, 769–776 (2001).

15 Cross, D. A., Alessi, D. R., Cohen, P., Andjelkovich, M. & Hemmings, B. A. Inhibition of glycogen synthase kinase-3 by insulin mediated by protein kinase B. Nature 378, 785–789 (1995).

16 Patel, P. & Woodgett, J. R. Glycogen Synthase Kinase 3: A Kinase for All Pathways? Curr Top Dev Biol 123, 277–302 (2017).

17 Hur, E. M. & Zhou, F. Q. GSK3 signalling in neural development. Nat Rev Neurosci 11, 539–551 (2010).

18 Tan, T. C. et al. Cdk5 is essential for synaptic vesicle endocytosis. Nat Cell Biol 5, 701–710 (2003).

19 Anggono, V. & Robinson, P. J. Syndapin I and endophilin I bind overlapping proline-rich regions of dynamin I: role in synaptic vesicle endocytosis. J Neurochem 102, 931–943 (2007).

20 Solomaha, E., Szeto, F. L., Yousef, M. A. & Palfrey, H. C. Kinetics of Src homology 3 domain association with the proline-rich domain of dynamins: specificity, occlusion, and the effects of phosphorylation. J Biol Chem 280, 23147–23156 (2005).

21 Cole, A. R. et al. Distinct priming kinases contribute to differential regulation of collapsin response mediator proteins by glycogen synthase kinase-3 in vivo. J Biol Chem 281, 16591–16598 (2006).

22 Wang, L. H. & Strittmatter, S. M. Brain CRMP forms heterotetramers similar to liver dihydropyrimidinase. J Neurochem 69, 2261–2269 (1997).

23 Deo, R. C. et al. Structural bases for CRMP function in plexin-dependent semaphorin3A signaling. EMBO J 23, 9–22 (2004).

24 Quach, T. T., Honnorat, J., Kolattukudy, P. E., Khanna, R. & Duchemin, A. M. CRMPs: critical molecules for neurite morphogenesis and neuropsychiatric diseases. Mol Psychiatry 20, 1037–1045 (2015).

25 Tanaka, H., Morimura, R. & Ohshima, T. Dpysl2 (CRMP2) and Dpysl3 (CRMP4) phosphorylation by Cdk5 and DYRK2 is required for proper positioning of Rohon-Beard neurons and neural crest cells during neurulation in zebrafish. Dev Biol 370, 223–236 (2012).

26 Yamashita, N. & Goshima, Y. Collapsin response mediator proteins regulate neuronal development and plasticity by switching their phosphorylation status. Mol Neurobiol 45, 234–246 (2012).

27 Karsan, A. et al. Quantitative proteomic analysis of sokotrasterol sulfate-stimulated primary human endothelial cells. Mol Cell Proteomics 4, 191–204 (2005).

28 Dickson, B. J. & Gilestro, G. F. Regulation of commissural axon pathfinding by slit and its Robo receptors. Annu Rev Cell Dev Biol 22, 651–675(2006).

29 Genet, G. et al. Endophilin-A2 dependent VEGFR2 endocytosis promotes sprouting angiogenesis. Nat Commun 10, 2350 (2019).

30 Jiang, H., Guo, W., Liang, X. & Rao, Y. Both the establishment and the maintenance of neuronal polarity require active mechanisms: critical roles of GSK-3beta and its upstream regulators. Cell 120, 123–135 (2005).

31 Byun, J. et al. Slit2 inactivates GSK3beta to signal neurite outgrowth inhibition. PLoS One 7, e51895 (2012).

32 Gao, F. J. et al. GSK-3beta Phosphorylation of Cytoplasmic Dynein Reduces Ndel1 Binding to Intermediate Chains and Alters Dynein Motility. Traffic 16, 941–961(2015).

33 Klinman, E., Tokito, M. & Holzbaur, E. L. F. CDK5-dependent activation of dynein in the axon initial segment regulates polarized cargo transport in neurons. Traffic 18, 808–824(2017).

34 Day, C. A. et al. Microtubule motors power plasma membrane tubulation in clathrin-independent endocytosis. Traffic 16, 572–590 (2015).

35 Renard, H. F. et al. Endophilin-A2 functions in membrane scission in clathrin-independent endocytosis. Nature 517, 493–496 (2015).

36 Firestone, A. J. et al. Small-molecule inhibitors of the AAA+ ATPase motor cytoplasmic dynein. Nature 484, 125–129 (2012).

37 Burkhardt, J. K., Echeverri, C. J., Nilsson, T. & Vallee, R. B. Overexpression of the dynamitin (p50) subunit of the dynactin complex disrupts dynein-dependent maintenance of membrane organelle distribution. J Cell Biol 139, 469–484 (1997).

38 Rietdorf, J. et al. Kinesin-dependent movement on microtubules precedes actin-based motility of vaccinia virus. Nat Cell Biol 3, 992–1000 (2001).

39 Prokic, I., Cowling, B. S. & Laporte, J. Amphiphysin 2 (BIN1) in physiology and diseases. J Mol Med (Berl) 92, 453–463 (2014).

40 Lichte, B. et al. Amphiphysin, a novel protein associated with synaptic vesicles. EMBO J 11, 2521–2530 (1992).

41 Sanchez-Barrena, M. J. et al. Bin2 is a membrane sculpting N-BAR protein that influences leucocyte podosomes, motility and phagocytosis. PLoS One 7, e52401(2012).

42 Ge, K. & Prendergast, G. C. Bin2, a functionally nonredundant member of the BAR adaptor gene family. Genomics 67, 210–220 (2000).

43 Ramjaun, A. R. & McPherson, P. S. Multiple Amphiphysin II Splice Variants Display Differential. J Neurochem. 7, 2369–2376 (1998).

44 Tomizawa, K. et al. Cophosphorylation of amphiphysin I and dynamin I by Cdk5 regulates clathrin-mediated endocytosis of synaptic vesicles. J Cell Biol 163, 813–824(2003).

45 Marks, B. & McMahon, H. T. Calcium triggers calcineurin-dependent synaptic vesicle recycling in mammalian nerve terminals. Curr Biol 8, 740–749 (1998).

46 Lai, M. M. et al. The calcineurin-dynamin 1 complex as a calcium sensor for synaptic vesicle endocytosis. J Biol Chem 274, 25963–25966 (1999).

47 Niethammer, M. et al. NUDEL Is a Novel Cdk5 Substrate that Associates with LIS1 and Cytoplasmic Dynein. Neuron 28, 697–711 (2000).

48 Fumoto, K., Hoogenraad, C. C. & Kikuchi, A. GSK-3beta-regulated interaction of BICD with dynein is involved in microtubule anchorage at centrosome. EMBO J 25, 5670–5682(2006).

49 Klinman, E. & Holzbaur, E. L. Stress-Induced CDK5 Activation Disrupts Axonal Transport via Lis1/Ndel1/Dynein. Cell Rep 12, 462–473 (2015).

50 Chapman, D. E. et al. Regulation of in vivo dynein force production by CDK5 and 14-3-3epsilon and KIAA0528. Nat Commun 10, 228 (2019).

51 Bianchi, M. et al. Regulation of FAK Ser-722 phosphorylation and kinase activity by GSK3 and PP1 during cell spreading and migration. Biochem J 391, 359–370(2005).

52 Inoki, K. et al. TSC2 integrates Wnt and energy signals via a coordinated phosphorylation by AMPK and GSK3 to regulate cell growth. Cell 126, 955–968 (2006).

53 Matta, S. et al. LRRK2 controls an EndoA phosphorylation cycle in synaptic endocytosis. Neuron 75, 1008–1021 (2012).

54 Murakami, N., Bolton, D. & Hwang, Y. W. Dyrk1A binds to multiple endocytic proteins required for formation of clathrin-coated vesicles. Biochemistry 48, 9297–9305(2009).

55 Wu, X., Gan, B., Yoo, Y. & Guan, J. L. FAK-mediated src phosphorylation of endophilin A2 inhibits endocytosis of MT1-MMP and promotes ECM degradation. Dev Cell 9, 185–196(2005).

56 Hoshina, S., Ueffing, M. & Weinstein, B. Growth factor-induced DNA synthesis in cells that overproduce protein kinase C. J Cell Physiol 145, 262–267 (1990).

57 Silva, J. C., Gorenstein, M. V., Li, G. Z., Vissers, J. P. & Geromanos, S. J. Absolute quantification of proteins by LCMSE: a virtue of parallel MS acquisition. Mol Cell Proteomics 5, 144–156 (2006).

