## Supplementary material for "Cdk5 and GSK3β inhibit Fast Endophilin-Mediated Endocytosis": Ferreira et al. Supplementary Figures

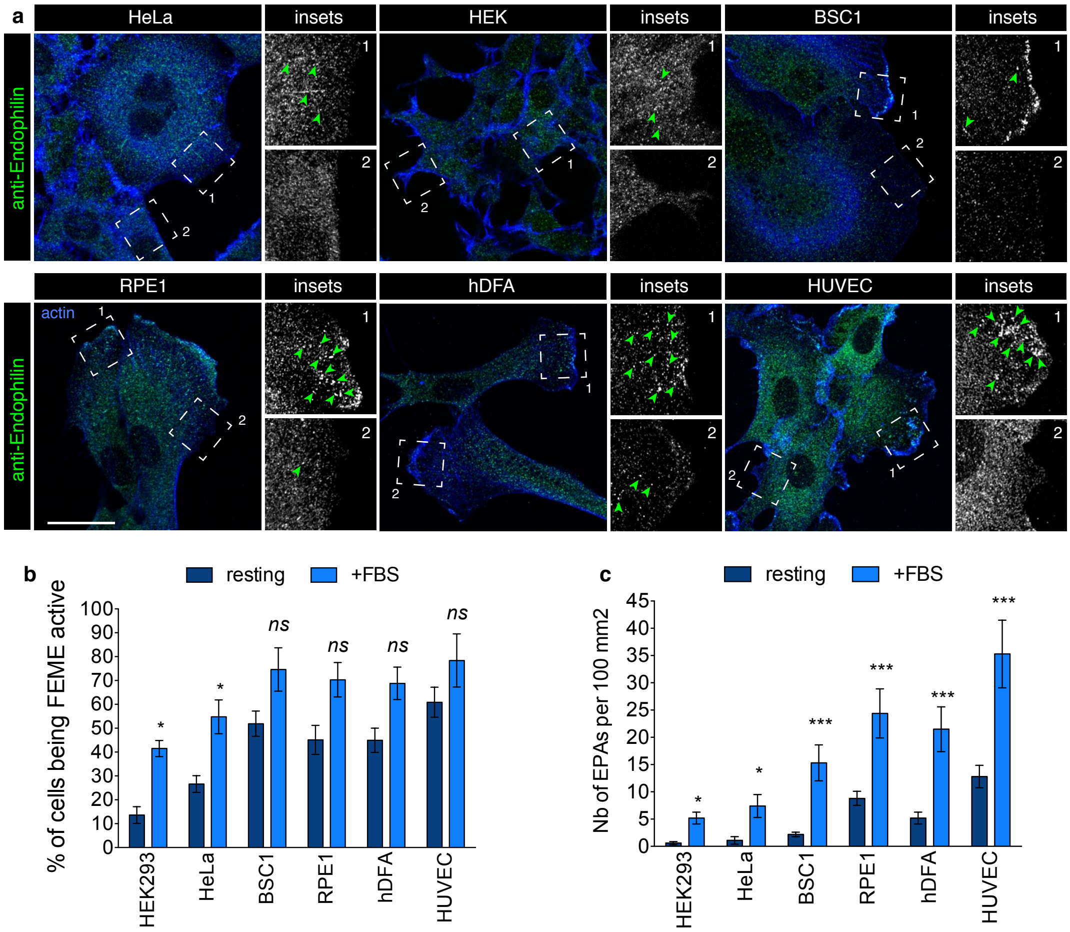


**Supplementary Figure 1**: related to Fig. 1 and 2

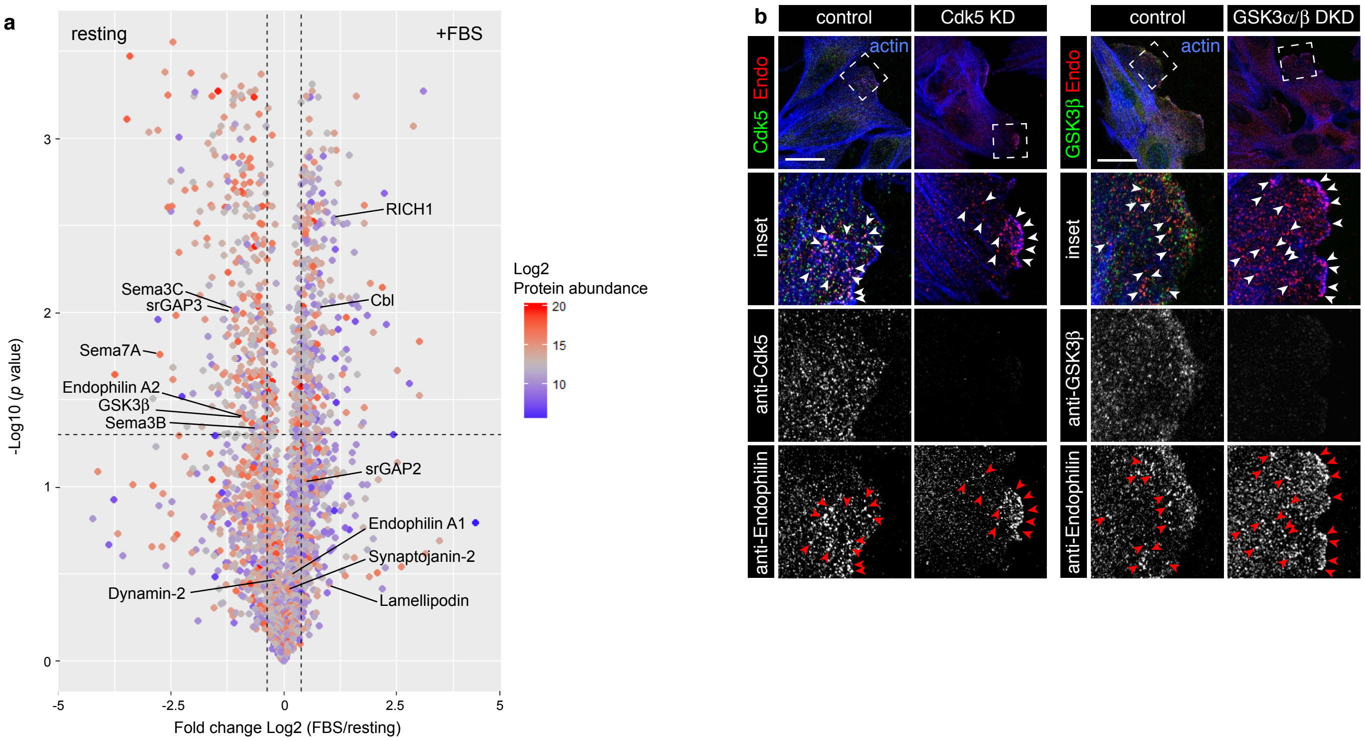


**Supplementary Figure 2**: related to Fig. 3

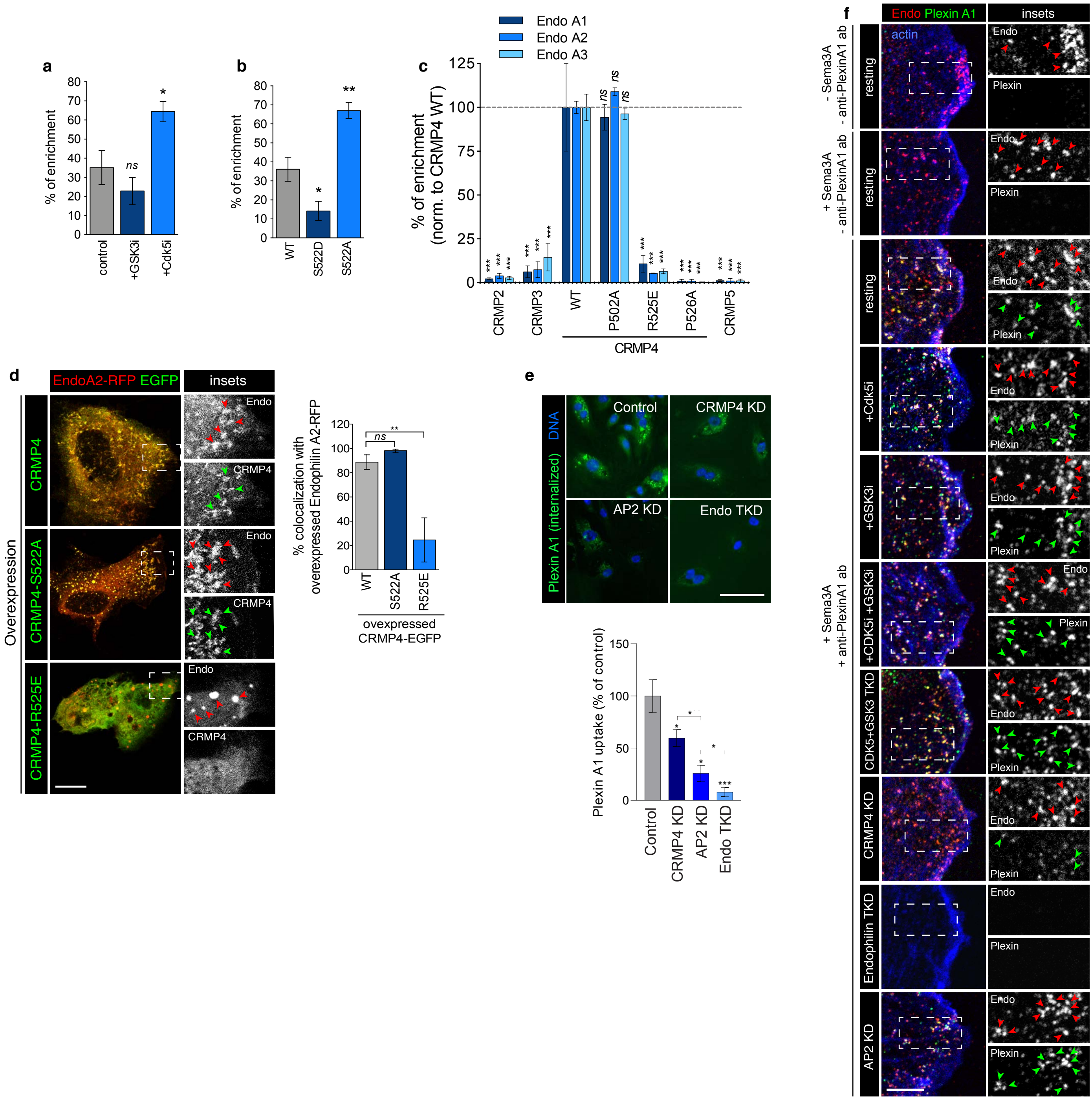


**Supplementary Figure 3**: related to Fig. 5

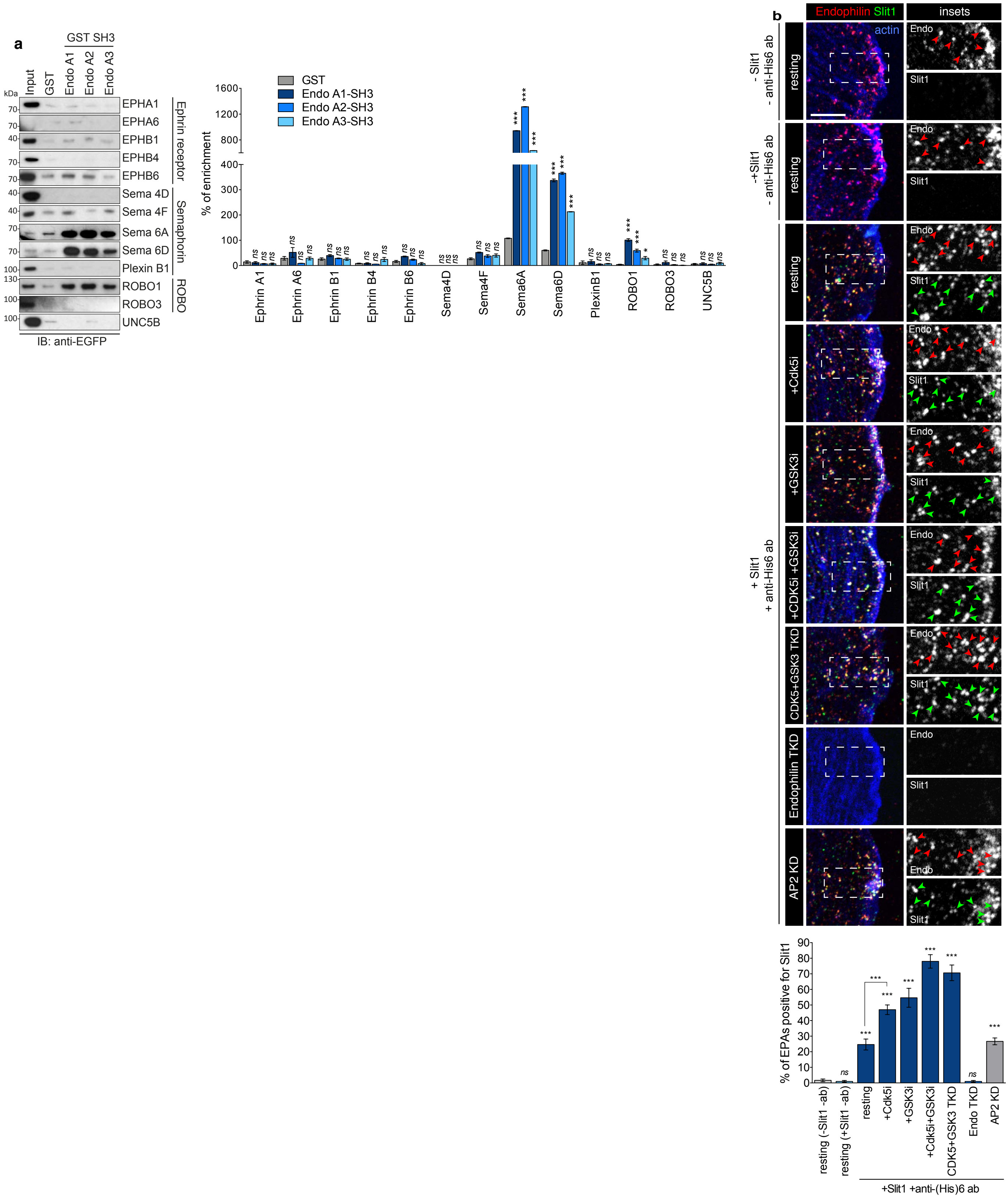

**Supplementary Figure 4**: related to Fig. 5

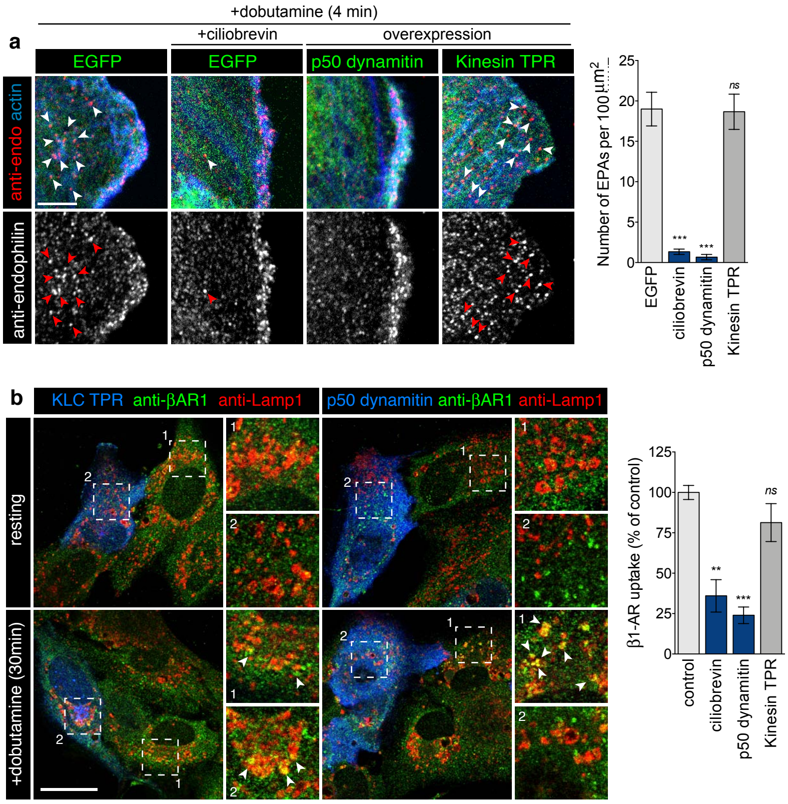


**Supplementary Figure 5**: related to Fig. 6

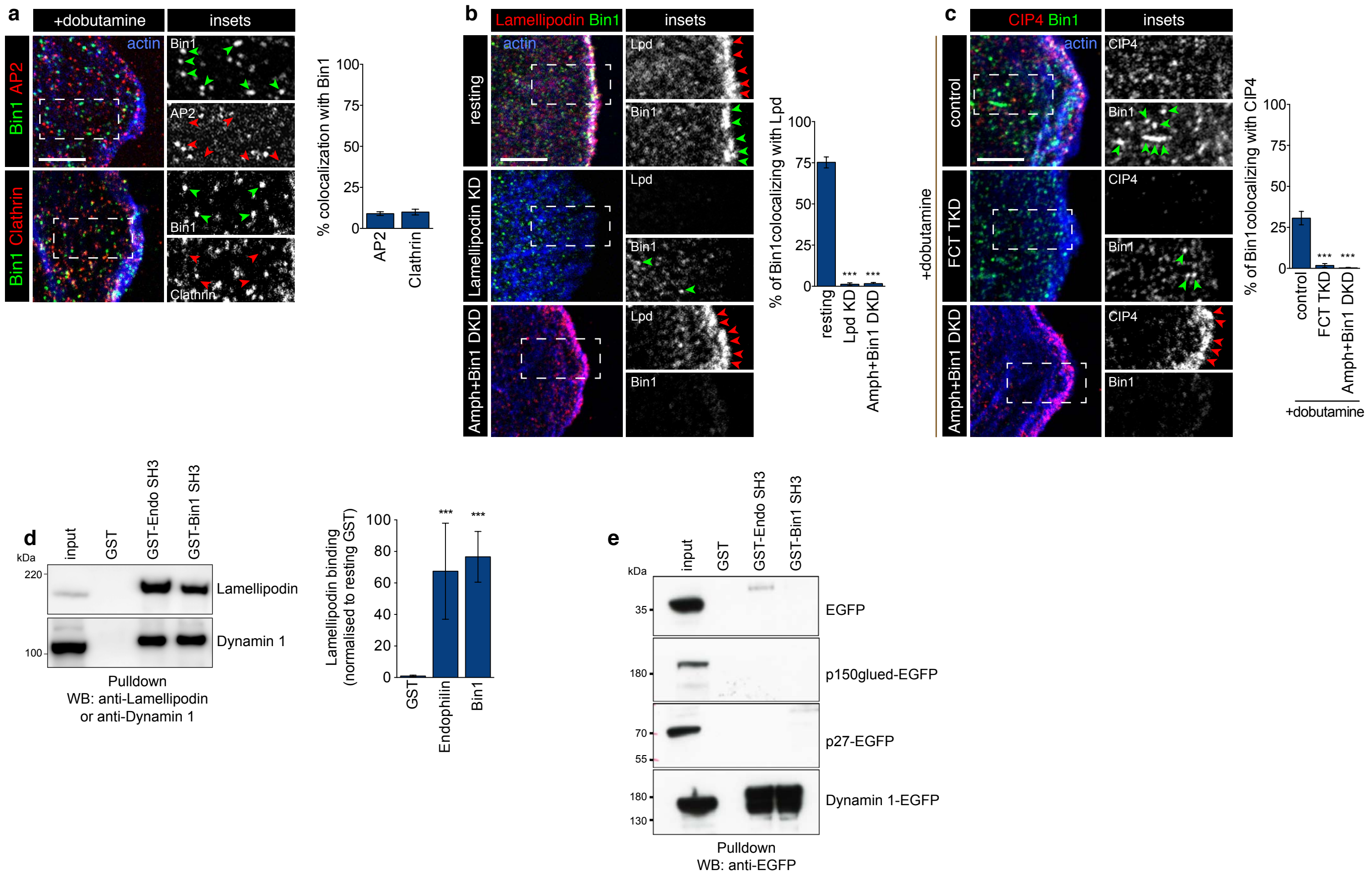


**Supplementary Figure 6**: related to Fig. 7
